# Perturbative formulation of general continuous-time Markov model of sequence evolution via insertions/deletions, Part III: Algorithm for first approximation

**DOI:** 10.1101/023614

**Authors:** Kiyoshi Ezawa, Dan Graur, Giddy Landan

## Abstract

**Background:** Insertions and deletions (indels) account for more nucleotide differences between two related DNA sequences than substitutions do, and thus it is imperative to develop a stochastic evolutionary model that enables us to reliably calculate the probability of the sequence evolution through indel processes. In a separate paper (Ezawa, Graur and Landan 2015a), we established an *ab initio* perturbative formulation of a continuous-time Markov model of the evolution of an *entire* sequence via insertions and deletions. And we showed that, under a certain set of conditions, the *ab initio* probability of an alignment can be factorized into the product of an overall factor and contributions from regions (or local alignments) separated by gapless columns. Moreover, in another separate paper (Ezawa, Graur and Landan 2015b), we performed concrete perturbation analyses on all types of local pairwise alignments (PWAs) and some typical types of local multiple sequence alignments (MSAs). The analyses indicated that even the fewest-indel terms alone can quite accurately approximate the probabilities of local alignments, as long as the segments and the branches in the tree are of modest lengths.

**Results:** To examine whether or not the fewest-indel terms alone can well approximate the alignment probabilities of more general types of local MSAs as well, and as a first step toward the automatic application of our *ab initio* perturbative formulation, we developed an algorithm that calculates the first approximation of the probability of a given MSA under a given parameter setting including a phylogenetic tree. The algorithm first chops the MSA into gapped and gapless segments, second enumerates all parsimonious indel histories potentially responsible for each gapped segment, and finally calculates their contributions to the MSA probability. We performed validation analyses using more than ten million local MSAs. The results indicated that even the first approximation can quite accurately estimate the probability of each local MSA, as long as the gaps and tree branches are at most moderately long.

**Conclusions:** The newly developed algorithm, called LOLIPOG, brought our *ab initio* perturbation formulation at least one step closer to a practically useful method to quite accurately calculate the probability of a MSA under a given biologically realistic parameter setting.

[This paper and three other papers (Ezawa, Graur and Landan 2015a,b,c) describe a series of our efforts to develop, apply, and extend the *ab initio* perturbative formulation of a general continuous-time Markov model of indels.]

**List of abbreviations:**

HMM: hidden Markov model
indel: insertion/deletion
LHS: local history set
MSA: multiple sequence alignment
PAS: preserved ancestral site
PWA: pairwise alignment

## Introduction

The evolution of DNA, RNA, and protein sequences is driven by mutations such as base substitutions, insertions and deletions (indels), recombination, and other genomic rearrangements (*e.g.*, Graur and Li 2000; Gascuel 2005; Lynch 2007). Thus far, analyses on substitutions have predominated in the field of molecular evolutionary study, in particular using the probabilistic (or likelihood) theory of substitutions that is now widely accepted (e.g., Felsenstein 1981, 2004; Yang 2006). However, some recent comparative genomic analyses have revealed that indels account for more base differences between the genomes of closely related species than substitutions (*e.g*., Britten 2002; Britten et al. 2003; Kent *et al*. 2003; The International Chimpanzee Chromosome 22 Consortium 2004; The Chimpanzee Sequencing and Analysis Consortium 2005). It is therefore imperative to develop a stochastic model that enables us to reliably calculate the probability of sequence evolution via mutations including insertions and deletions.

Since the groundbreaking works by Bishop and Thompson (1986) and by Thorne, Kishino and Felsenstein (1991), there have been many efforts to calculate the alignment probabilities under the probabilistic models aiming to incorporate the effects of indels. Over the past few decades, such methods have greatly improved in terms of the computational efficiency and the scope of application (see, *e.g*., Rivas 2005; Bradley and Holmes 2007; Miklós et al. 2009). However, these methods, mostly based on hidden Markov models (HMMs) or transducer theories, have two fundamental problems, one regarding the theoretical grounds and the other regarding the biological realism. (See the “background” section in part I (Ezawa, Graur and Landan 2015a) for more details on these problems.)

To solve these two problems, we chose to base our study on an indel evolutionary model that is devoid of the problems from the beginning. The model we chose were a *genuine* stochastic evolutionary model, more specifically, a general continuous-time Markov model of the evolution of an *entire* sequence via indels along the time-axis. The model allows any indel rate parameters including length distributions, but it does not impose any unnatural restrictions on indels. In part I of this series of study (Ezawa, Graur and Landan 2015a), we established an *ab initio* perturbative formulation of the general continuous-time Markov model. We showed that, when the indel rate parameters satisfy a certain set of conditions, the *ab initio* probability of an alignment can be factorized into the product of an overall factor and contributions from regions (or local alignments) separated by gapless columns. In part II (Ezawa, Graur and Landan 2015b), we concretely calculated the fewest-indel contributions and the next-fewest-indel contributions to the probability of each local alignment, among all types of local pairwise alignments (PWAs) and some typical types of local multiple sequence alignments (MSAs). Our perturbation analyses indicated that even the fewest-indel contribution can approximate the probability of each local alignment quite accurately, as long as the local alignment is not so long and the branch lengths are at most moderately long. We are confident that this conclusion should be quite general on the local PWAs, because we exhausted all possible types of homology structures (Lunter et al. 2005). However, in order to claim that the conclusion holds generally also on local MSAs, we need a more extensive analysis, by exploring most of the local MSA patterns we could encounter in practical evolutionary processes.

For this purpose, in this study, we developed an algorithm to calculate such a “first-approximate” probability for an input MSA, under a given parameter setting including a phylogenetic tree. To validate our algorithm and the conclusion in part II, we conducted some simulation analyses. Using a *genuine* molecular evolution simulator, Dawg (Cartwright 2005), we created more than ten million local MSAs and counted the absolute frequency of, as well as the relative frequencies of ancestral states for, each local gap configuration. We used these frequencies as the “correct answers” to be compared to the first-approximate probabilities calculated only from the contributions by the fewest-indel histories. The results indicated that the conclusion in part II seems to hold for a more general set of local MSAs, and thus they demonstrated the use of the first-approximate probabilities under modest settings.

In Results, we describe the results of our validation analyses. In Discussion, we will discuss some possible improvements and applications of our theory and algorithm. The topics include the risks associated with the naïve application of our algorithm to *reconstructed* alignments. The Methods section details the algorithms and analyses. Subsection M1 of Methods describes our algorithm to calculate the first approximation of the probability of a given MSA. Subsection M2 of Methods describes the details on our validation analyses.

This paper is part III of a series of our papers that documents our efforts to develop, apply, and extend the *ab initio* perturbative formulation of the general continuous-time Markov model of sequence evolution via indels. Part I (Ezawa, Graur and Landan 2015a) gives the theoretical basis of this entire study. Part II (Ezawa, Graur and Landan 2015b) describes concrete perturbation calculations and examines the applicable ranges of other probabilistic models of indels. Part III (this paper) describes our algorithm to calculate the first approximation of the probability of a given MSA and simulation analyses to validate the algorithm. Finally, part IV (Ezawa, Graur and Landan 2015c) discusses how our formulation can incorporate substitutions and other mutations, such as duplications and inversions.

This paper basically uses the same conventions as used in part I (Ezawa, Graur and Landan 2015a). See its Section 2 for details if necessary. And, as in part I, the following terminology is used. The term “an indel process” means a series of successive indel events with both the order and the specific timings specified, and the term “an indel history” means a series of successive indel events with only the order specified. And, throughout this paper, the union symbol, such as in *A* ∪*B* and 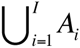, should be regarded as the union of *mutually disjoint* sets (*i.e.,* those satisfying *A* ∩ *B* = ∅ and *A*_*i*_ ∩ *A*_*j*_ = ∅ for *i* ≠ *j* (∈ {1,…, *I*}), respectively, where ∅ is an empty set), unless otherwise stated.

## Results

In Subsection 1.2 of part II (Ezawa, Graur and Landan 2015b), we saw that, as long as the indel lengths and the branch lengths are at most moderate, the contributions from the fewest-indel histories alone can well approximate the multiplication factors for any local gap configurations in PWAs. And, in Subsection 1.3 of part II, we saw that this is also the case with some typical gap configurations in MSAs. In MSAs, however, there could be many patterns of gap configurations, in addition to those examined in Subsection 1.3 of part II. Thus, to examine whether or not the contributions by the fewest-indel histories can in general well approximate the multiplication factors for local gap-configurations of MSAs, we conducted simulation analyses.

First, we developed an algorithm that performs the following series of three processes (Figure 1 A). (i) It first partitions a given MSA into an alternating series of gapped and gapless segments. (ii) It second enumerates the fewest-indel local histories (*i.e.*, the parsimonious local indel histories) giving rise to each of the gapped segments. And (iii) it third calculates the “fewest-indel approximation” of the multiplication factor for each gapped segment (Eq.(1.1.2a) of part II) by summing the contributions from all the fewest-indel local histories. The absolute probability of the given MSA is approximated by the product of the probability, 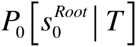 (given by Eq.(4.2.9b) of part I (Ezawa, Graur and Landan 2015a)), that a reference root state 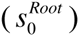 is kept throughout the tree *T*, and the approximate multiplication factors for the gapped segments (calculated in parts I & II). (Because the algorithm only enumerates the fewest-indel histories, it ignores “null local indel histories” that leave no traces in the MSA, which were discussed in Subsection 3.3 of part I.) As a by-product, the algorithm also calculates the relative probabilities among the fewest-indel local histories that can give rise to each gapped segment. For details of the algorithm, see Methods M1 and Figures 1-6. The algorithm is currently implemented only under Dawg’s indel model (Cartwright 2005; see also Eqs.(2.4.4a,b,c) of part I), and the indel length distributions can be chosen from power-law and geometric distributions. We provided the current implementation of the algorithm in a prototype package named LOLIPOG (<u>lo</u>g-<u>li</u>kelihood for the <u>p</u>attern <u>o</u>f <u>g</u>aps), which we made available at the FTP repository of the Bioinformatics Organization (Ezawa 2013).

**Figure 1.**
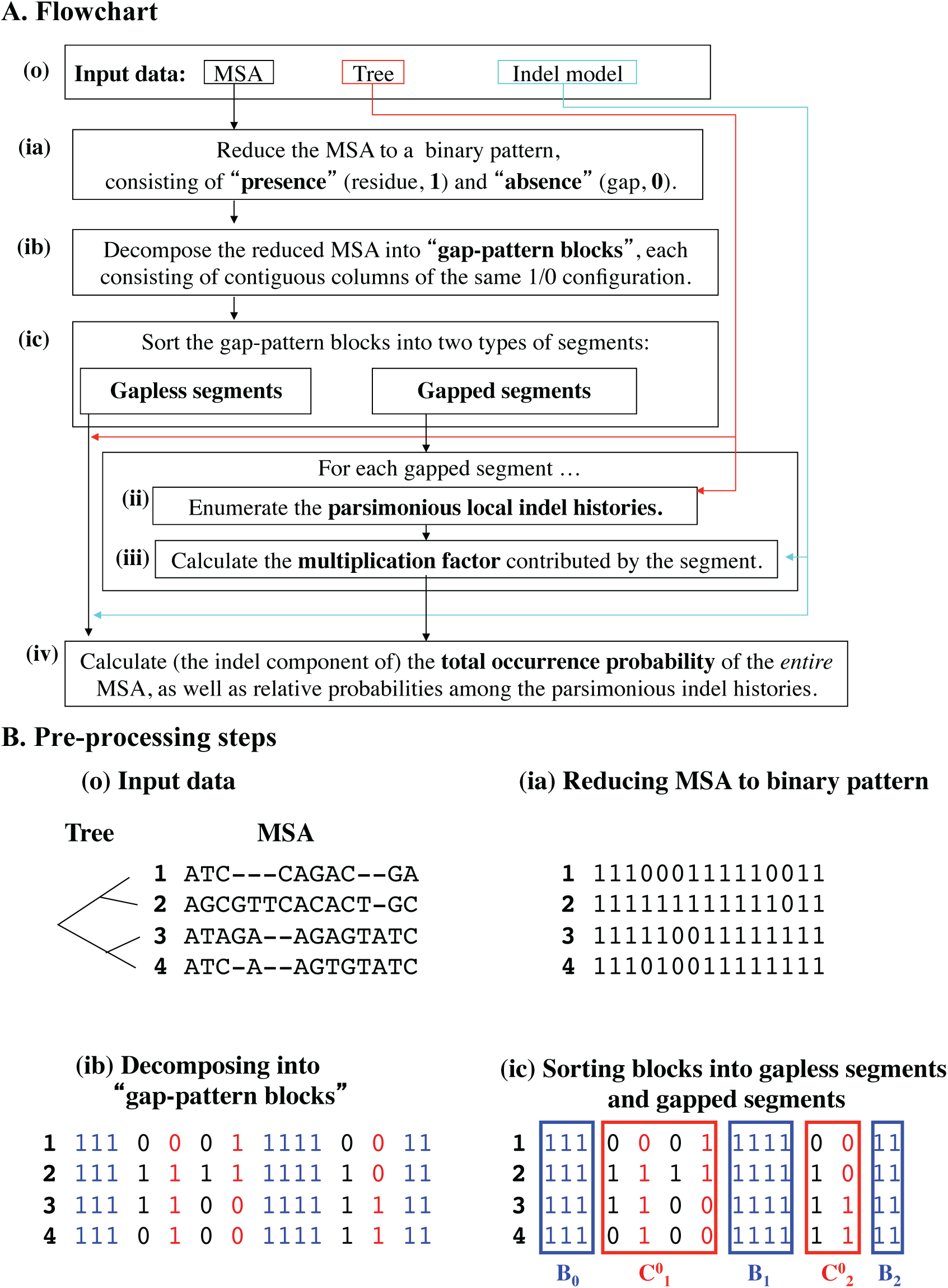
Overall workflow in our algorithm to calculate the MSA probability. The entire algorithm consists of steps (ia), (ib), (ic), (ii) and (iii), processing the input (o) into the final output at step (iv). **(A)** The flowchart. **(B)** The schematic illustration of the pre-processing steps (ia-ic). The input data [ **(o)**] consists mainly of a MSA (of DNA sequences here) and a phylogenetic tree of the aligned sequences (labeled with boldface numbers). An evolutionary model via indels is assumed to be given but is omitted here. Step **(ia)** reduces the input MSA to a binary 1/0 pattern, with 1 and 0 representing the “presence” (of a residue) and the “absence” (*i.e*., a gap), respectively. Step **(ib)** decomposes the binary pattern into “gap-pattern block"s, or “block”s for short, each of which consists of contiguous columns of a given 1/0 pattern. Here each block is represented as a rectangular array of neighboring cells with a particular color. Step **(ic)** sorts the blocks into gapless segments (each represented as contiguous blue cells enclosed by a blue rectangle labeled *B*_*k*_ (with *k* = 0,1, 2 )) and gapped segments *K* (each represented as contiguous cells enclosed by a red rectangle labeled 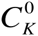 (with *K* = 1, 2 )). See M1.1 (in Methods) for more details. [NOTE: The set of all gapped segments, 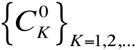, is a subset of 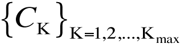, which is the set of all regions that can accommodate local indel histories along the tree.]

**Figure 2.**
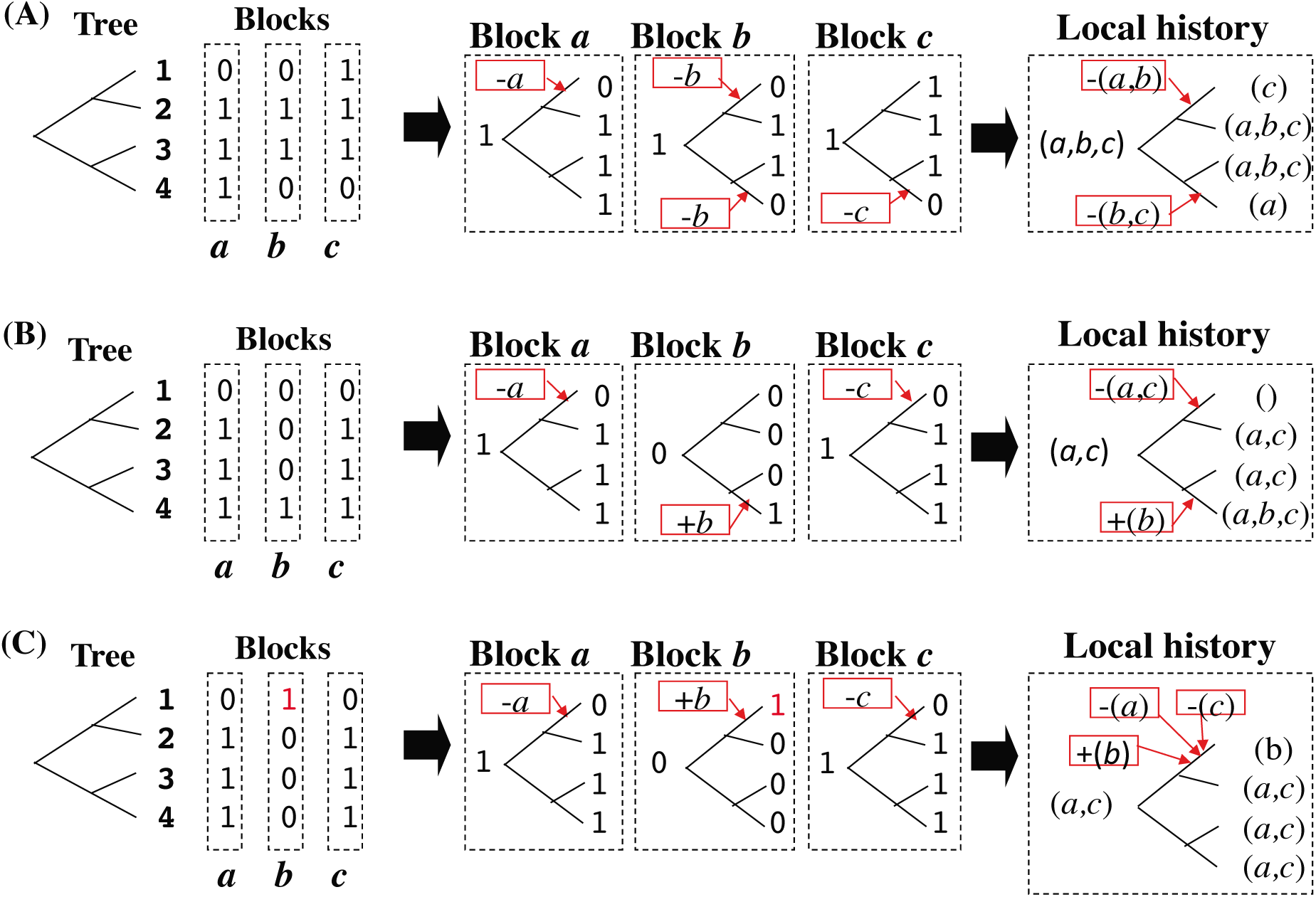
Merging indel events in effectively contiguous gap-pattern blocks. In each panel, given a gapped segment consisting of contiguous gap-pattern blocks (“block”s), and a phylogenetic tree of aligned sequences (left), the Dollo parsimonious history for each block is first inferred (middle), then the indel histories in the effectively contiguous blocks are merged if they are of the same type and occur along the same branch (right). As in Figure 1 B, a “1” and a “0” represent the presence state (*i.e.,* a residue) and the absence state (*i.e.*, a gap), respectively. Note that each column under the “Blocks” (left) represents a gap-pattern block, and not necessarily a single column, in the MSA. In the indel histories in the middle step, “+x” and “-y” represent the insertion of block “x” and the deletion of block “y”, respectively. In the local indel histories in the final step (on the right), blocks in the same parentheses after the “+” or the “-” sign, respectively, are inserted or deleted simultaneously. **(A)** Merging indel events in literally contiguous blocks. **(B)** Merging indel events in two blocks separated by a (run of) block(s) in which no downstream nodes with the “presence” state interrupt the merger. **(C)** In this case, the deletions of block *a* and block *c*, both along the exterior branch leading to sequence 1, cannot be merged because they are interrupted by the downstream node with the “presence” state (the red “1”) in block *b*.

**Figure 3.**
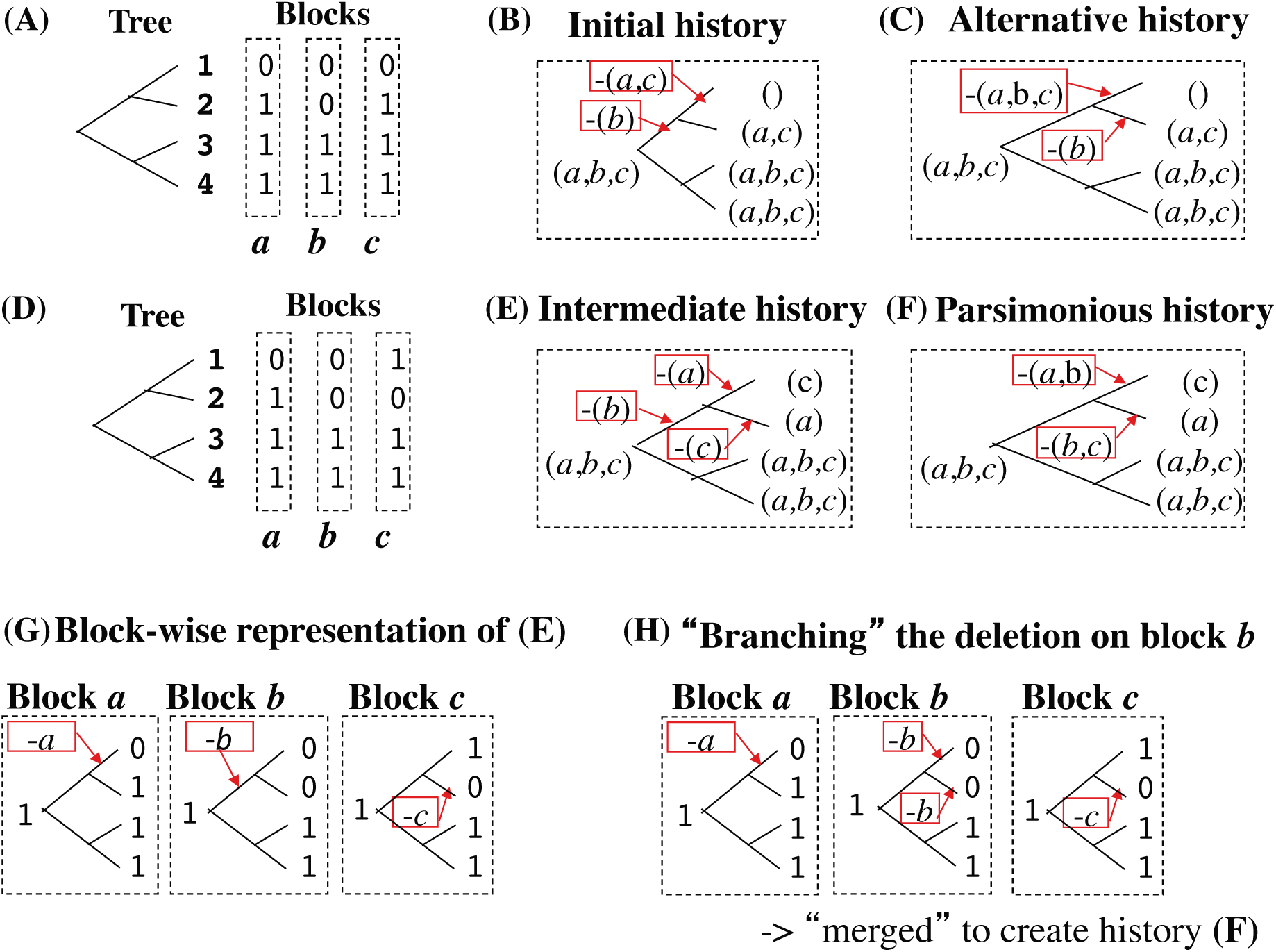
Looking for parsimonious local indel histories. For the gap-configuration (under the “Blocks”) and the tree shown in **(A)**, the initial step infers the history in **(B)**, but there is actually another parsimonious history **(C)**. For the segment and the tree shown in **(D)**, using the history in **(E)** as an “intermediate” point always reachable from the initial history, we can find the actual parsimonious history shown in **(F)**. **(G,H)** a “branch-and-merge” operation performed on the situation in **(D)**. **(G)** Looking closely at the indel history in (E), we see that a deletion of a subsequence in block *b* occurs along the branch of the common ancestor of sequences 1 and 2. With this history as a starting point, in the “branching” step **(H)**, the deletion is re-interpreted as deletions along the child branches. Finally, merging the resulting deletions with the effectively contiguous deletion(s) gives the local indel history in **(F)** in this example.

**Figure 4.**
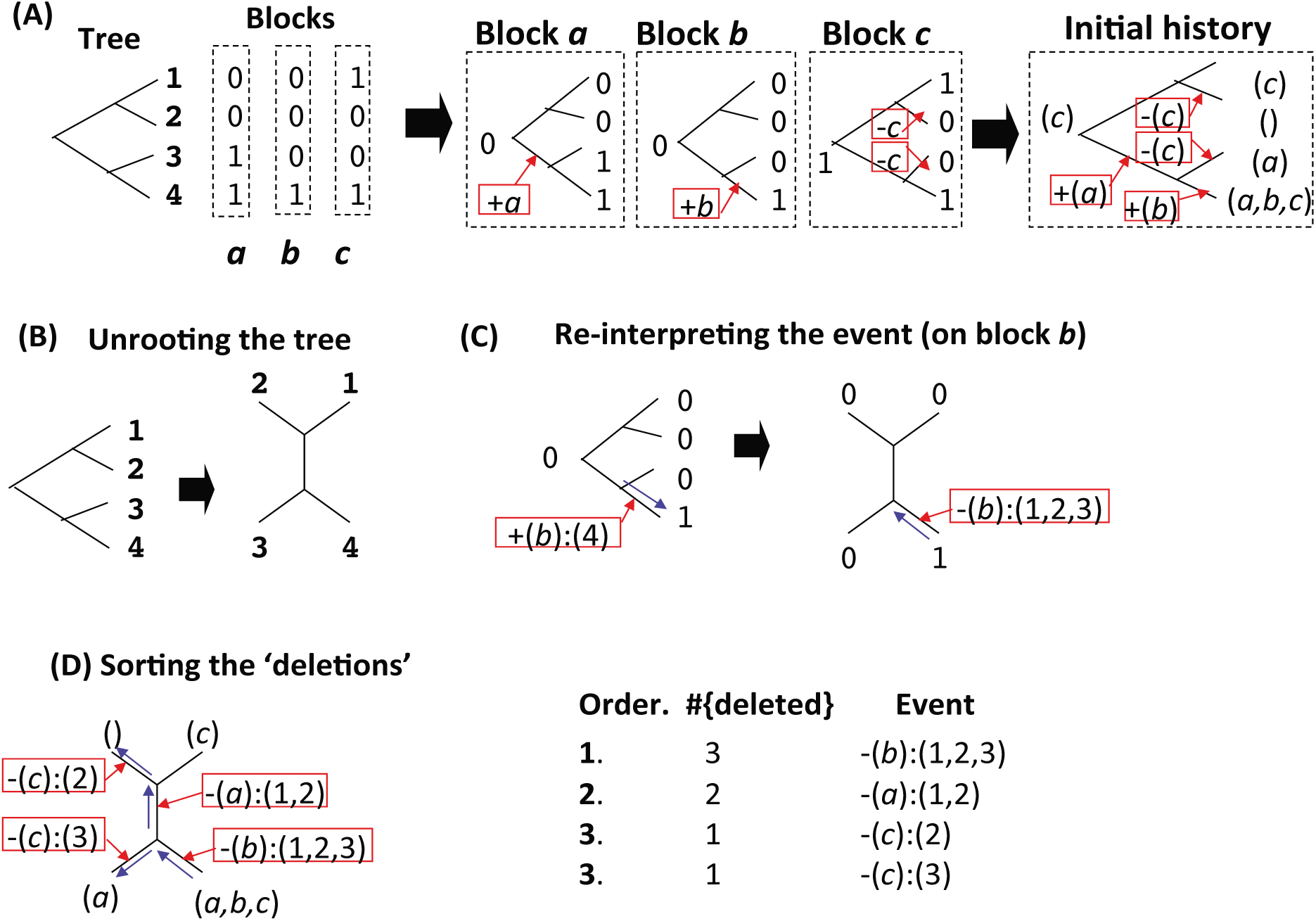
Sorting indel events that will undergo “branch-and-merge” processes. **(A)** The initial local indel history (right), given a gapped segment (under the “Blocks”) and a sequence tree (left). **(B)** If the input tree is rooted, it gets unrooted. **(C)** Then, an insertion event (as in block *b* in this example) can be re-interpreted as a ‘deletion’ event by reversing the (virtual) time direction (represented by a blue arrow). Here, “+(*b*):(4)” denotes that block *b* was inserted into sequence 4, and “-(b):(1,2,3)” denotes that block *b* was deleted from the (‘last common ancestor’ of) sequences 1, 2, and 3. Similarly, “+(a):(3,4)” in the original history will also be re-interpreted as “- (a):(1,2).” **(D)** In this way, we can re-interpret all the indel events as ‘deletions’ (left), and sort them in descending order of the number of ‘deleted’ sequences (right).

**Figure 5.**
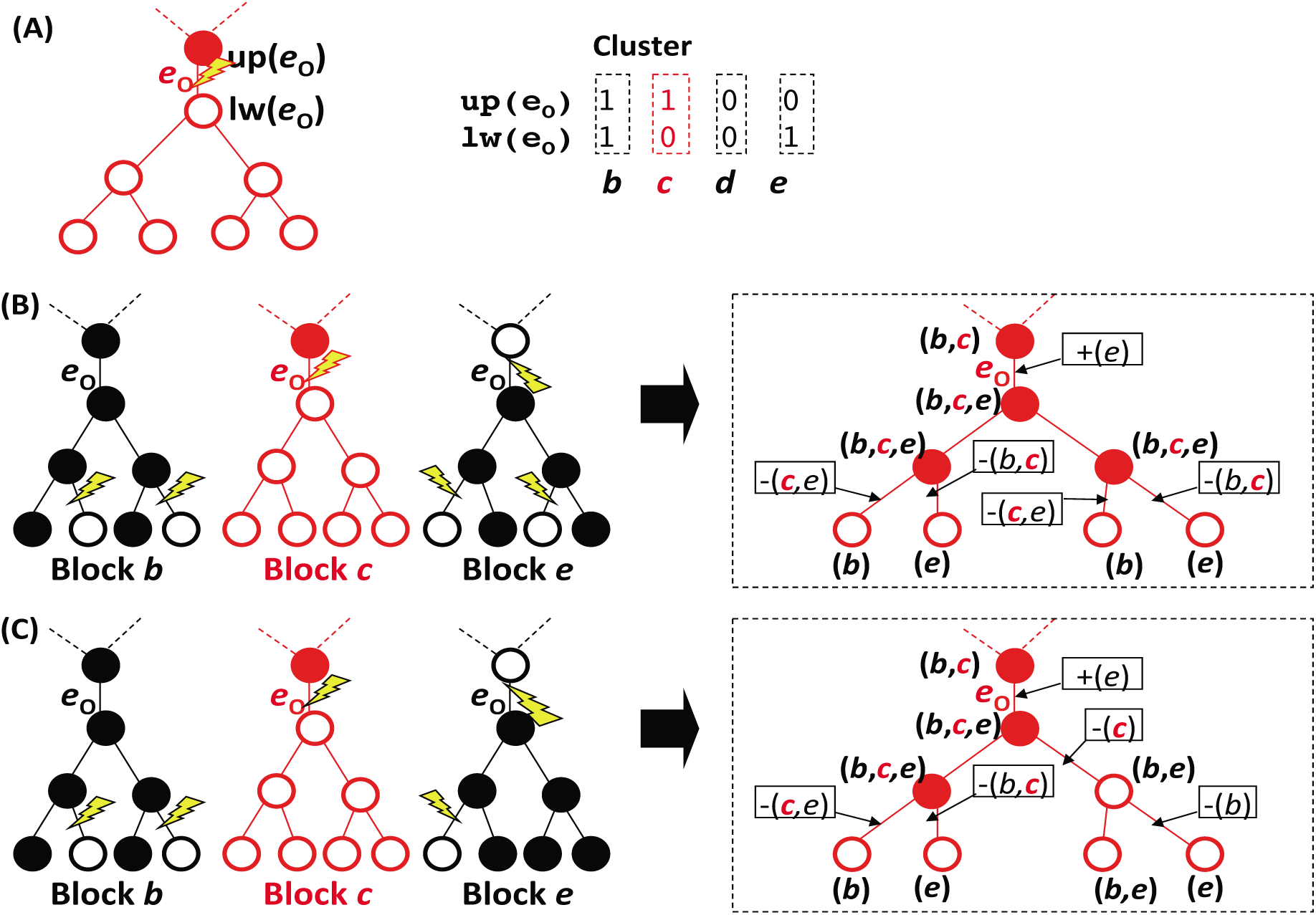
Composite “branch-and-merge” operation: schematic illustrations. Panel **(A)** partially shows an input local indel history. The red tree on the left shows the “presence” and “absence” states (the solid and open red circles, respectively), as well as a single ‘deletion’ event (yellow lightening bolt) along branch *e*_*o*_, on gappattern block *c*. The symbols *up*(*e*_*o*_) and *lw*(*e*_*o*_) label the nodes at the ‘upper-end’ and the ‘lower-end,’ respectively, of branch *e*_*o*_. The dashed lines ‘above’ *up*(*e*_*o*_) are the remaining part of the tree, whose details don’t matter here. The array of “0” and “1” on the right briefly represents the pairwise alignment of the sequences at nodes *up*(*e*_*o*_) and *lw*(*e*_*o*_ ). We assume that gap-pattern block *c* (in red), which concerns us the most, is effectively flanked by blocks *b* and *e*. The block *d* was skipped because it has the “absence” state at nodes *up*(*e*_*o*_) and *lw*(*e*_*o*_), as well as at all nodes in the ‘downstream’ of them (not shown). **(B, C)** Shown on the left are the input indel history on block *c* and those on the effectively flanking blocks *b* and *e*. On the right is the local indel history on the entire segment consisting of blocks *b*, *c* and *e*, superimposed by the history on block *c* (in red). Both of the histories are after a composite “branch-and-merge” operation on the ‘deletion’ of block *c* along branch *e*_*o*_. At each node, (*w*,…,*z*) represents that blocks *w*,…,*z* are “present.” Along each branch, +(*x*,…, *y*) denotes that blocks *x*,…, *y* are ‘inserted,’ and −(*x*,…, *y*) denotes that blocks *x*,…, *y* are ‘deleted.’ **(B)** The ‘deletions’ involving effectively flanking blocks, *b* and *e,* along branches ‘under’ branch *e*_*o*_ (black-contoured lightening bolts), can ‘delete’ all the sequences ‘under’ *e*_*o*_ (left). In this case, the total number of indels reduces by 1 after the “branch-and-merge” operation (right), providing a local history that truly replaces the old candidate histories. **(C)** In this case, to ‘delete’ all the sequences ‘under’ branch *e*_*o*_, the ‘deletions’ involving effectively flanking blocks, *b* and *e*, are not enough (left). Actually, an additional ‘deletion’ is necessary (“-(c)” along a branch of the tree in the right dashed box). Thus, in this case, the total number of indels does not change after the “branch-and-merge” operation (right), and the resulting local history joins the set of current candidate histories.

**Figure 6.**
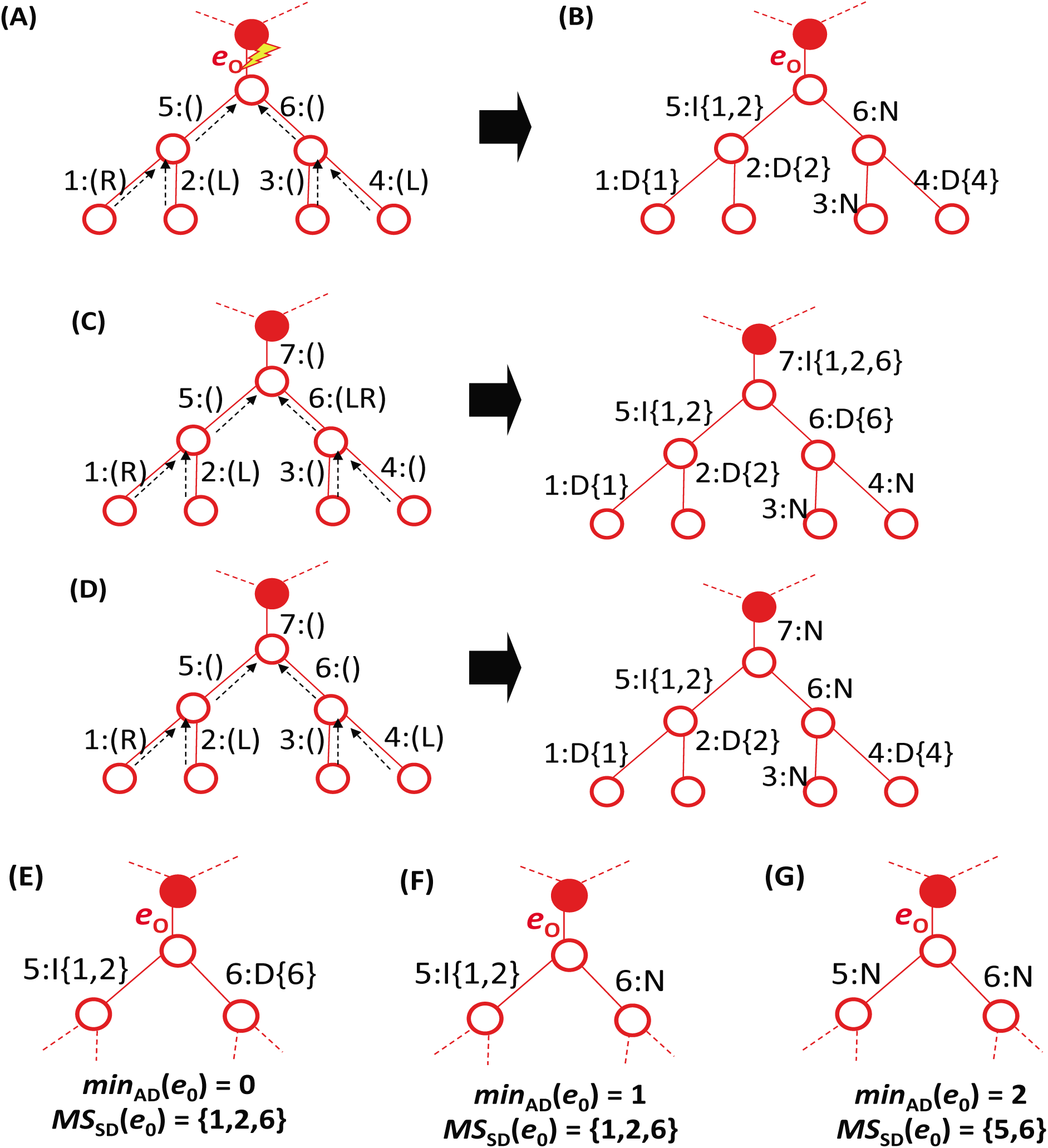
‘Bottom-up’ algorithm to search for minimal set of ‘deletion’ events in composite “branch-and-merge” operation. This schematic illustration uses the input local indel history on the left of panel (C) in Figure 5 as an example. **(A)** The algorithm starts from the external branches ‘under’ branch *e*_*o*_, and goes upwards until it reaches *e*_*o*_ (upward dashed arrows). Here, the branches are numbered from 1 to 6 (in order of being processed), to facilitate the explanation. Each branch is assigned a flanking ‘deletion’ status, “R”, “L”, “RL”, or “” (nothing), in parentheses after a colon, indicating that the branch undergoes (a) ‘deletion(s)’ involving the right-“flanking” block (*b in* Figure 5), the left-“flanking” block (*e in* Figure 5), both blocks, or none, respectively. **(B)** Annotating the branches. Each branch is classified as “directly absorbable” (D), if it itself is *not* assigned the status “” (branches 1, 2, and 4 in this example); it is classified as “indirectly absorbable” (I), if it is assigned the status “” and all of its child branches are classified as “directly” or “indirectly” “absorbable” (branch 5); or otherwise, it is classified as “non-absorbable” (N) (branches 3 and 6). The set of numbers in braces after the D/I symbol on each branch represents the minimal set of directly absorbable branches the ‘deletions’ along which can ‘delete’ *all* the sequences ‘under’ the branch in question. Branch 7 in panel **C** gives a more complex example of an “indirectly absorbable” branch. Branch 7 in panel **D** is a more complex example of a “non-absorbable” branch. **(E,F,G)** The algorithm finishes on branch *e*_*o*_ by giving the minimum number of additional ‘deletion’ events (min_*AD*_ (*e*_*o*_)), as well as the minimum set of ‘deletions’ that jointly substitute for the original ‘deletion’ (*MS*_*SD*_ (*e*_*o*_ )). **(E)** When all ‘child’ branches are (“directly” or “indirectly”) “absorbable.” **(F)** When some ‘child’ branches are “absorbable” and others are “non-absorbable”. **(G)** When all ‘child’ branches are “non-absorbable".

Second, to validate the component of the algorithm that enumerates all fewest-indel local histories potentially resulting in the gap configuration of a given gapped segment, we applied it to a set of simple MSAs each accompanied by a phylogenetic tree of the sequences (Methods M2.1 and Figures 7-27). The MSAs and accompanying trees were chosen to extensively cover typical cases of the gap-configurations of the segments and local indel histories that can generate them. We manually confirmed that our implementation of the algorithm certainly enumerates all conceivable fewest-indel histories that can generate each of the gap-configurations, except for some complex cases that are expected to be very rare (Methods M2.1; see also Discussion).

**Figure 7.**
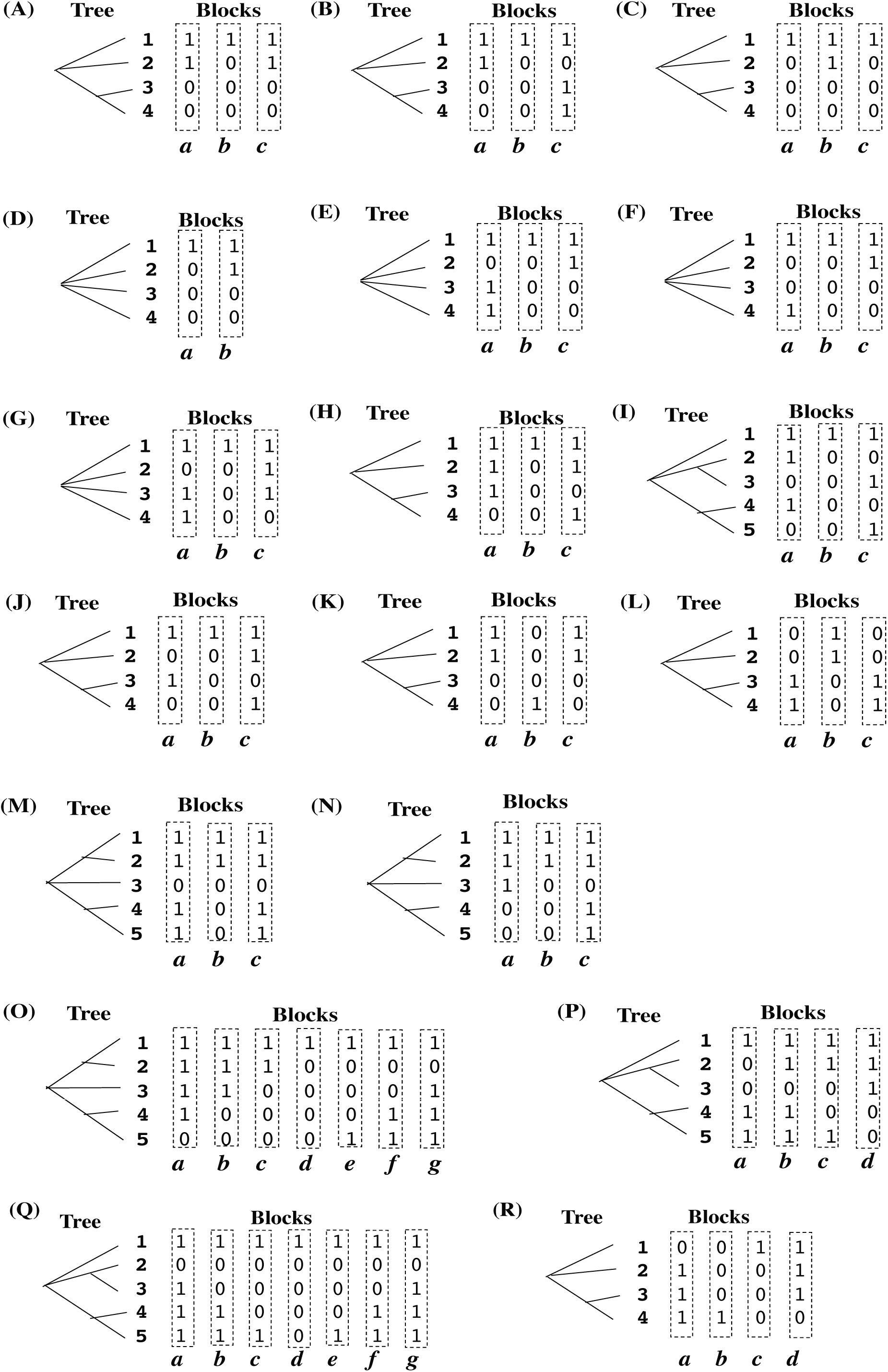
Input data for validation of our indel parsimony algorithm. Each of the panels, A through R, shows a two-component set of input data. One component is the gap-configuration of a gapped segment in a MSA, consisting of contiguous gap-pattern blocks (columns of “0”s denoting gaps and/or “1” s denoting residues, each enclosed by a dashed rectangle labeled with a bold italic alphabet). The other is a phylogenetic tree of aligned sequences (labeled with bold Arabic numerals).

**Figure 8.**
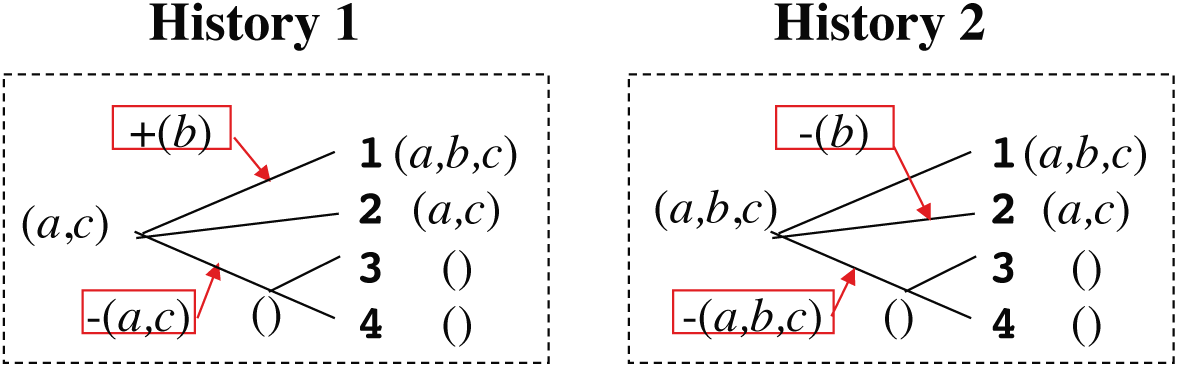
Local indel histories output by our parsimony algorithm, given input in Figure 7 A. The output is schematically illustrated using the phylogenetic tree of the aligned sequences. At each external node (labeled with a bold Arabic numeral), or at each internal node (unlabeled), “(*x,‥,z*)” denotes a set of blocks that are “present” in the corresponding (existing or ancestral) sequence. In particular, the “()” at a node represents the situation where all relevant blocks are “absent” from the corresponding sequence. In a red box, “+(*s,…,t*)” denotes an insertion of the subsequence consisting of blocks *s*,…, and *t*, and “-(*u,…,v*)” denotes a deletion of the subsequence consisting of blocks *u*,…, and *v*. A red arrow points to the branch along which the indel event occurred. (Nominal) time is supposed to run from left to right.

**Figure 9.**
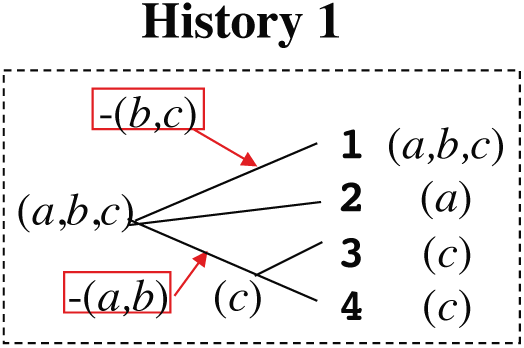
Local indel history output by our parsimony algorithm, given input in Figure 7 B. Notation is the same as in Figure 8.

**Figure 10.**
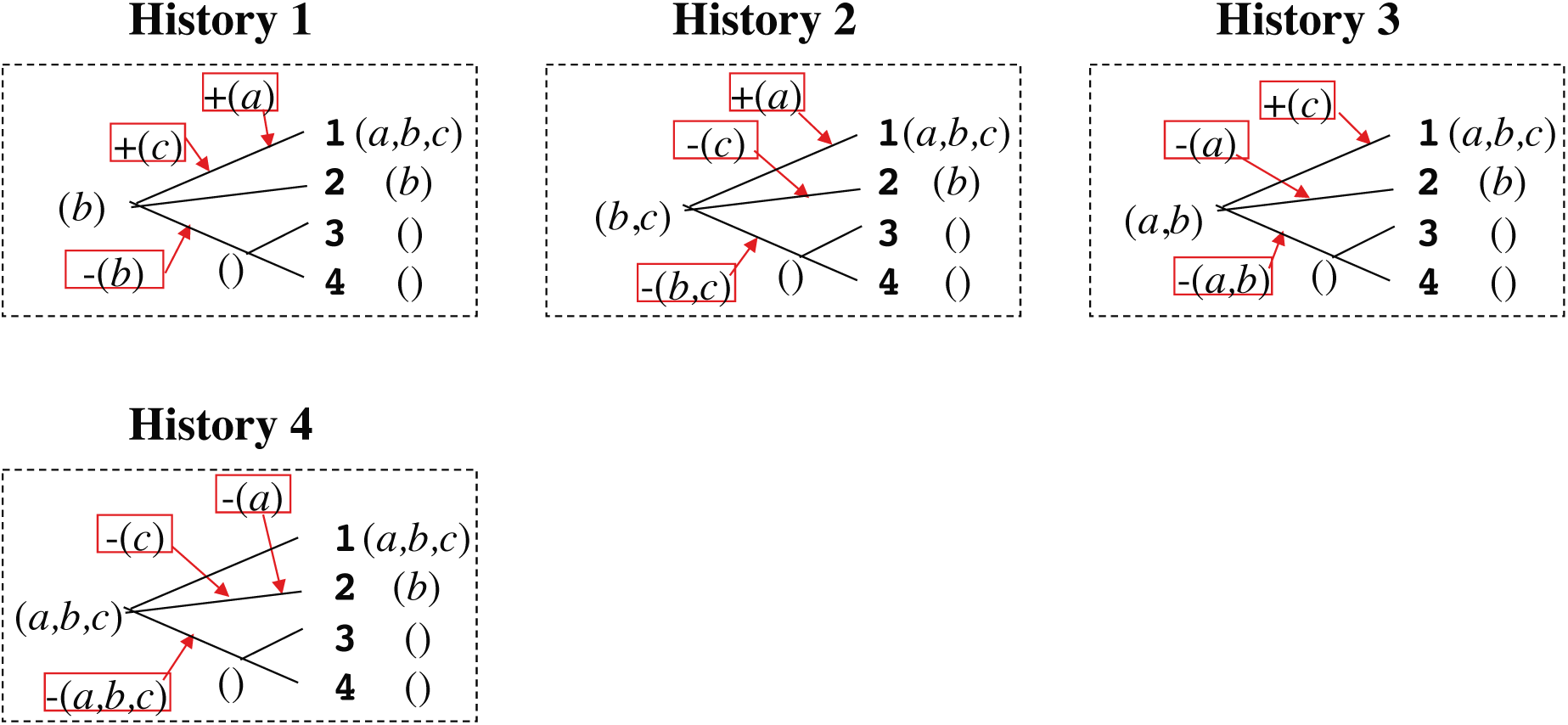
Local indel histories output by our parsimony algorithm, given input in Figure 7 C. Notation is the same as in Figure 8.

**Figure 11.**
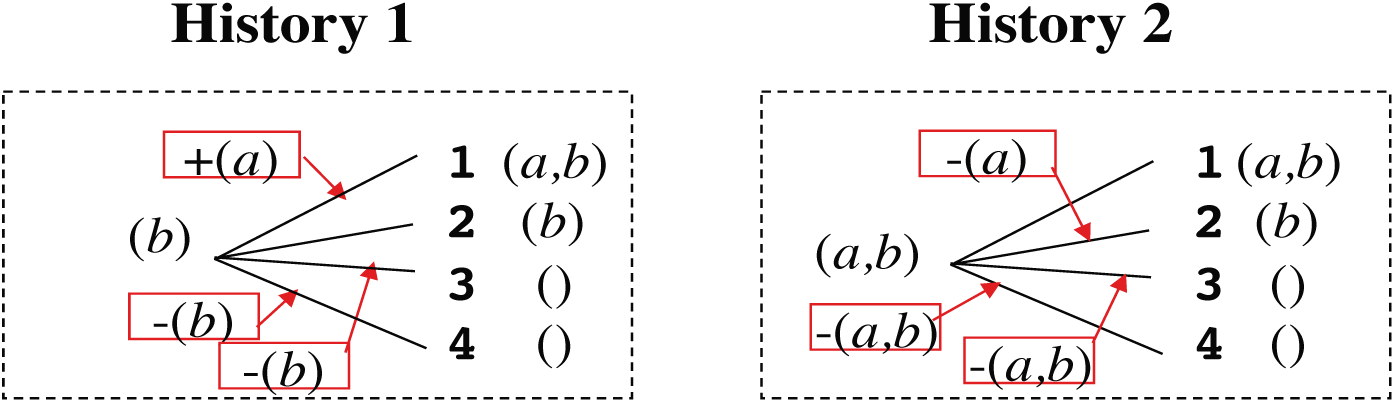
Local indel histories output by our parsimony algorithm, given input in Figure 7 D. Notation is the same as in Figure 8.

**Figure 12.**
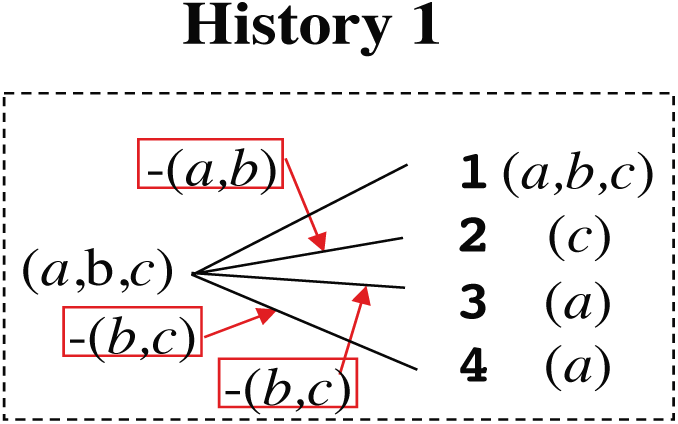
Local indel history output by our parsimony algorithm, given input in Figure 7 E. Notation is the same as in Figure 8.

**Figure 13.**
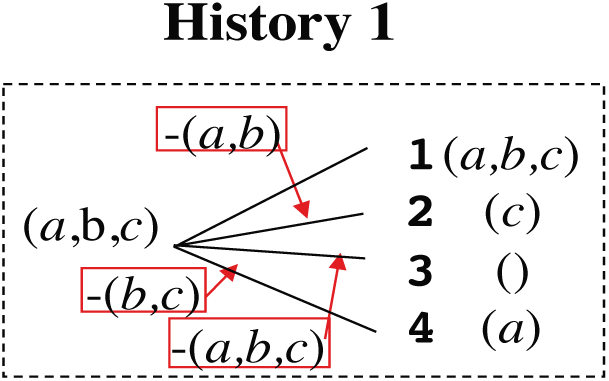
Local indel history output by our parsimony algorithm, given input in Figure 7 F. Notation is the same as in Figure 8.

**Figure 14.**
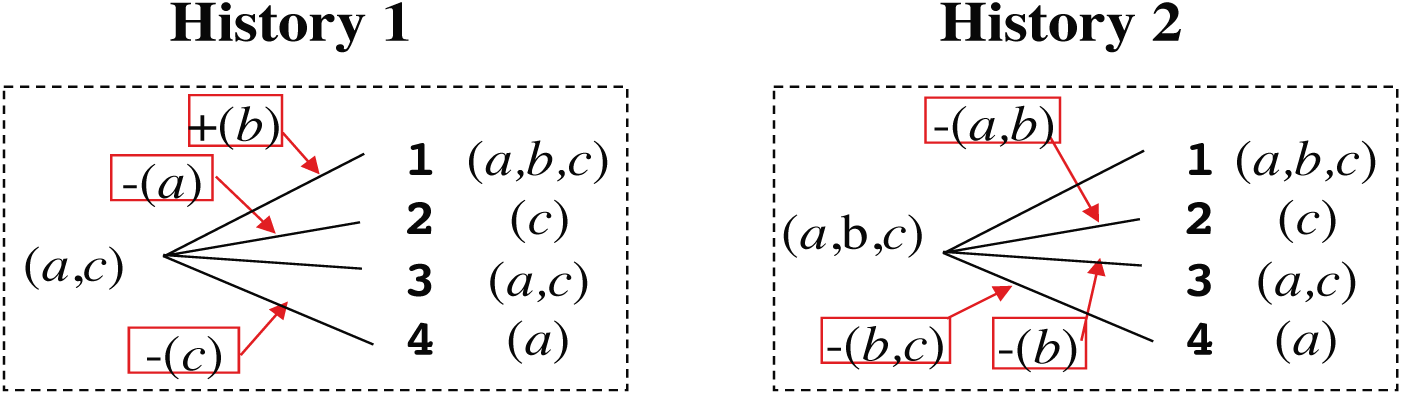
Local indel histories output by our parsimony algorithm, given input in Figure 7 G. Notation is the same as in Figure 8.

**Figure 15.**
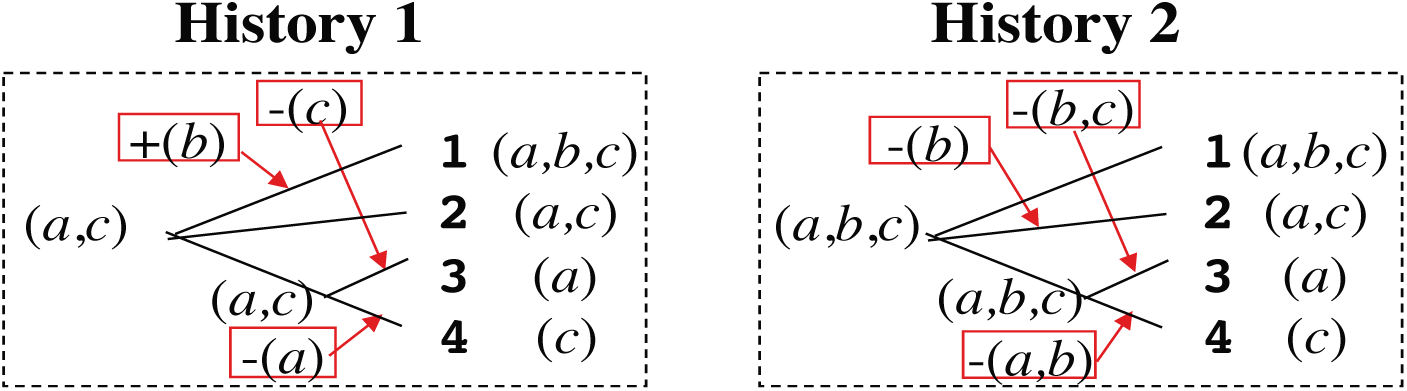
Local indel histories output by our parsimony algorithm, given input in Figure 7 H. Notation is the same as in Figure 8.

**Figure 16.**
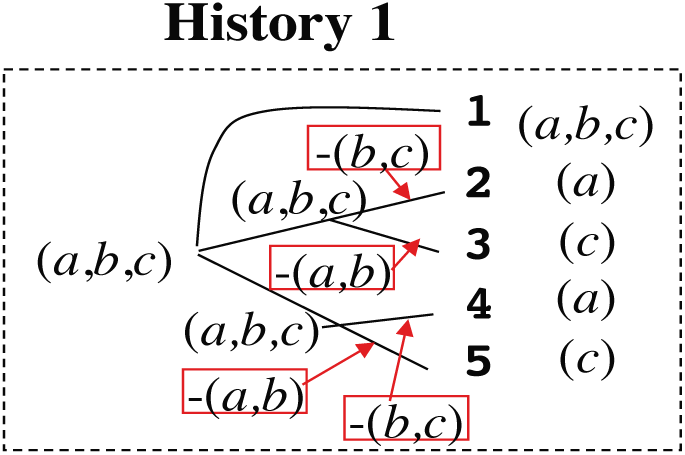
Local indel history output by our parsimony algorithm, given input in Figure 7 I. Notation is the same as in Figure 8.

**Figure 17.**
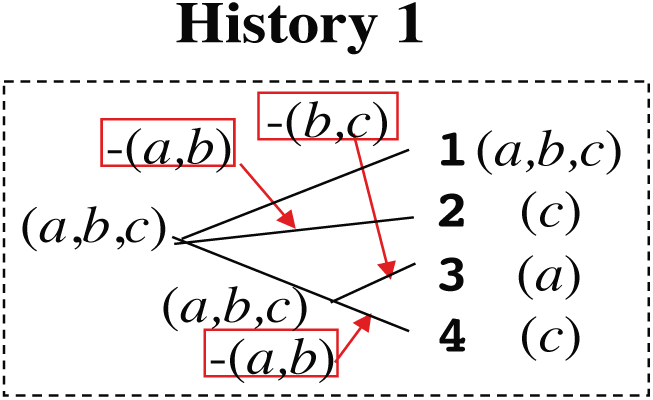
Local indel history output by our parsimony algorithm, given input in Figure 7 J. Notation is the same as in Figure 8.

**Figure 18.**
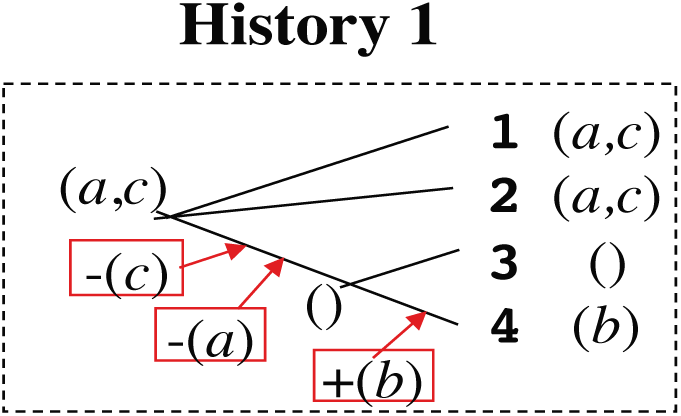
Local indel history output by our parsimony algorithm, given input in Figure 7 K. Notation is the same as in Figure 8.

**Figure 19.**
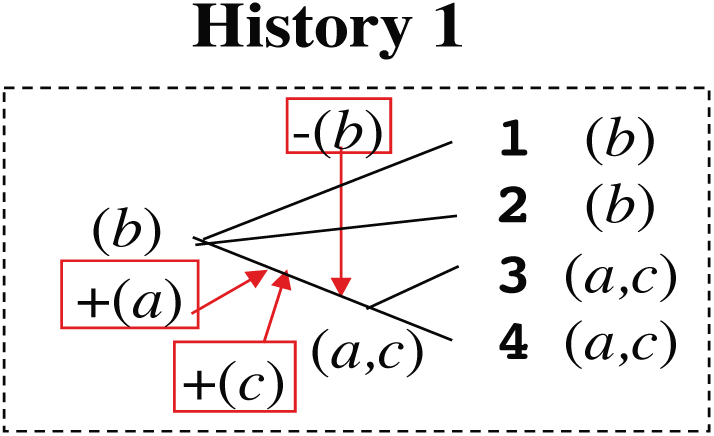
Local indel history output by our parsimony algorithm, given input in Figure 7 L. Notation is the same as in Figure 8. The temporal order of the indel events is changeable within some restrictions specific to the aligner or simulator.

**Figure 20.**
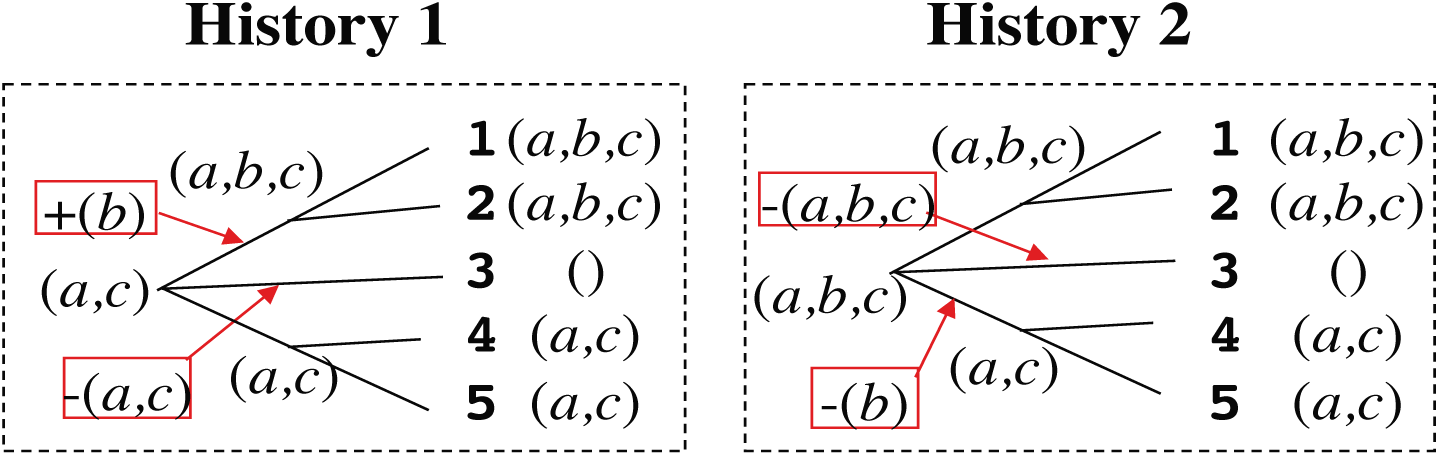
Local indel histories output by our parsimony algorithm, given input in Figure 7 M. Notation is the same as in Figure 8.

**Figure 21.**
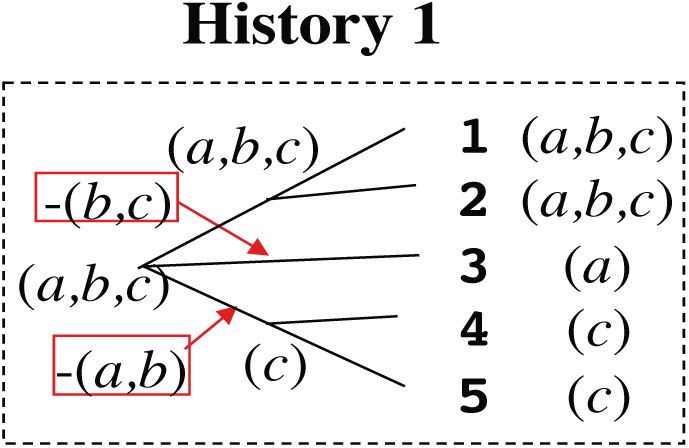
Local indel history output by our parsimony algorithm, given input in Figure 7 N. Notation is the same as in Figure 8.

**Figure 22.**
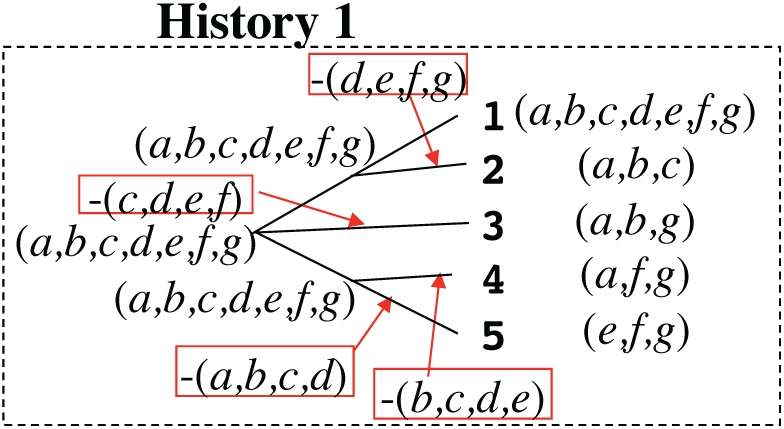
Local indel history output by our parsimony algorithm, given input in Figure 7 O. Notation is the same as in Figure 8.

**Figure 23.**
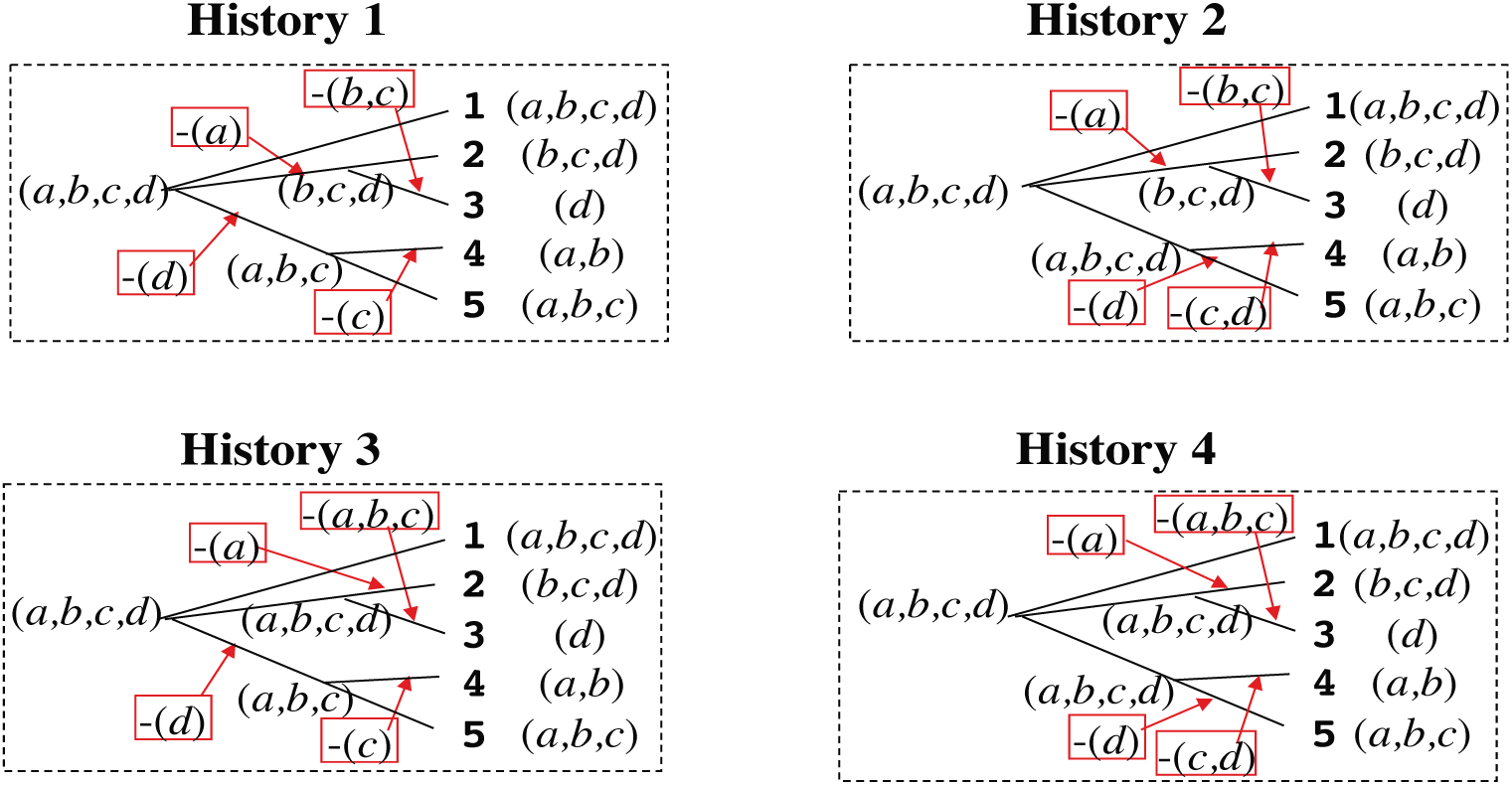
Local indel histories output by our parsimony algorithm, given input in Figure 7 P. Notation is the same as in Figure 8.

**Figure 24.**
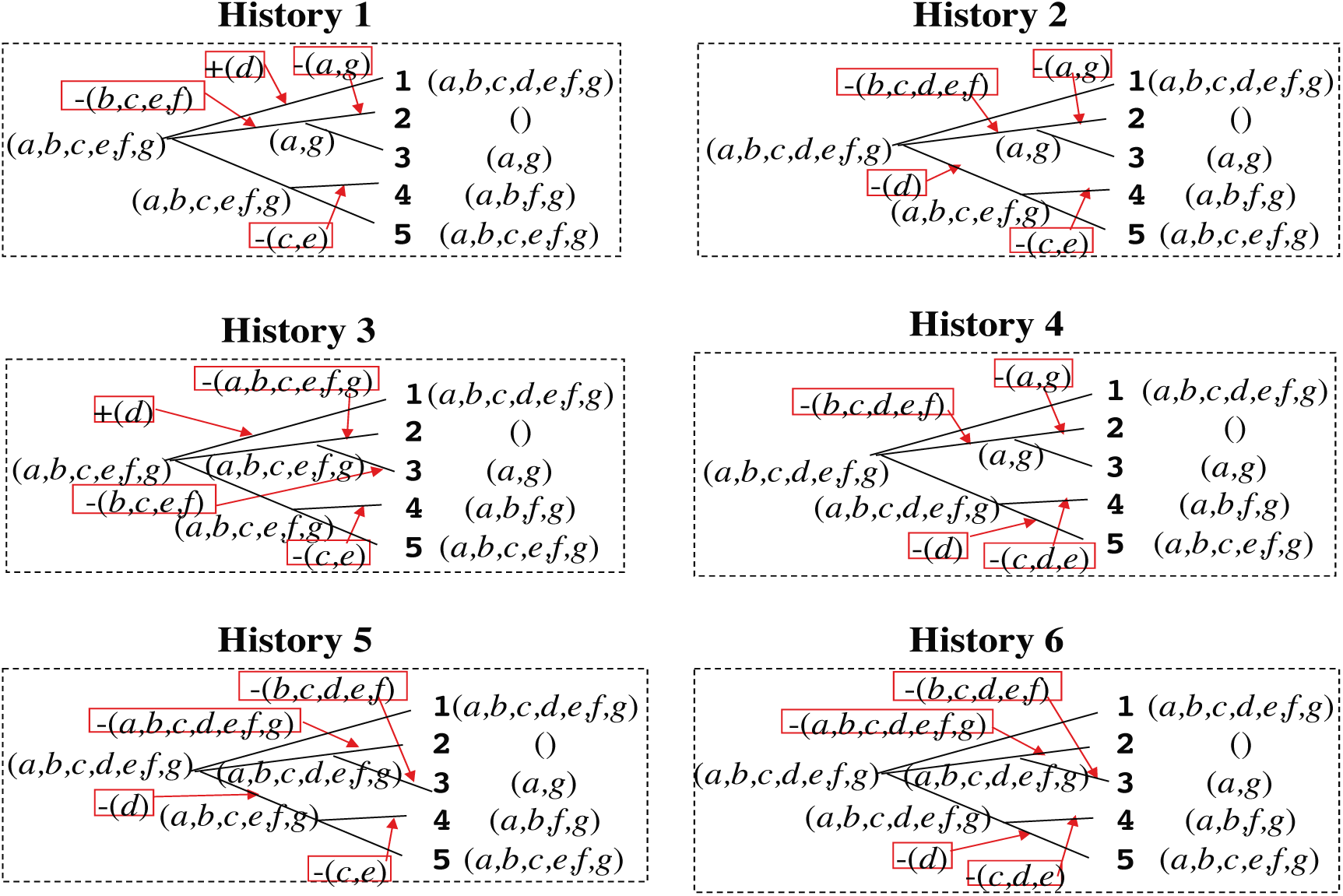
Local indel histories output by our parsimony algorithm, given input in Figure 7 Q. Notation is the same as in Figure 8.

**Figure 25.**
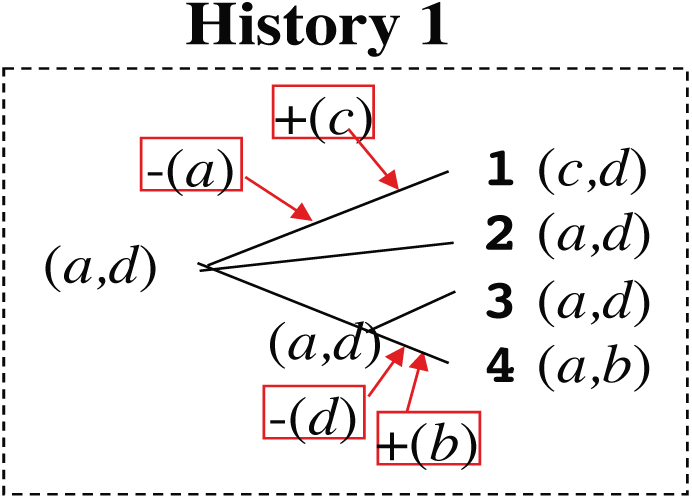
Local indel history output by our current parsimony algorithm, given input in Figure 7 R. Notation is the same as in Figure 8.

**Figure 26.**
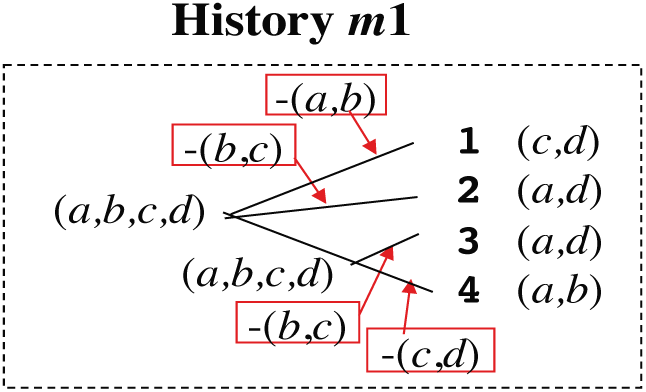
Typical local indel history that current implementation of our parsimony algorithm cannot find. This is a parsimonious local indel history that gives rise to the gapped segment as in panel R of Figure 7. See Methods M2.1 for details.

**Figure 27.**
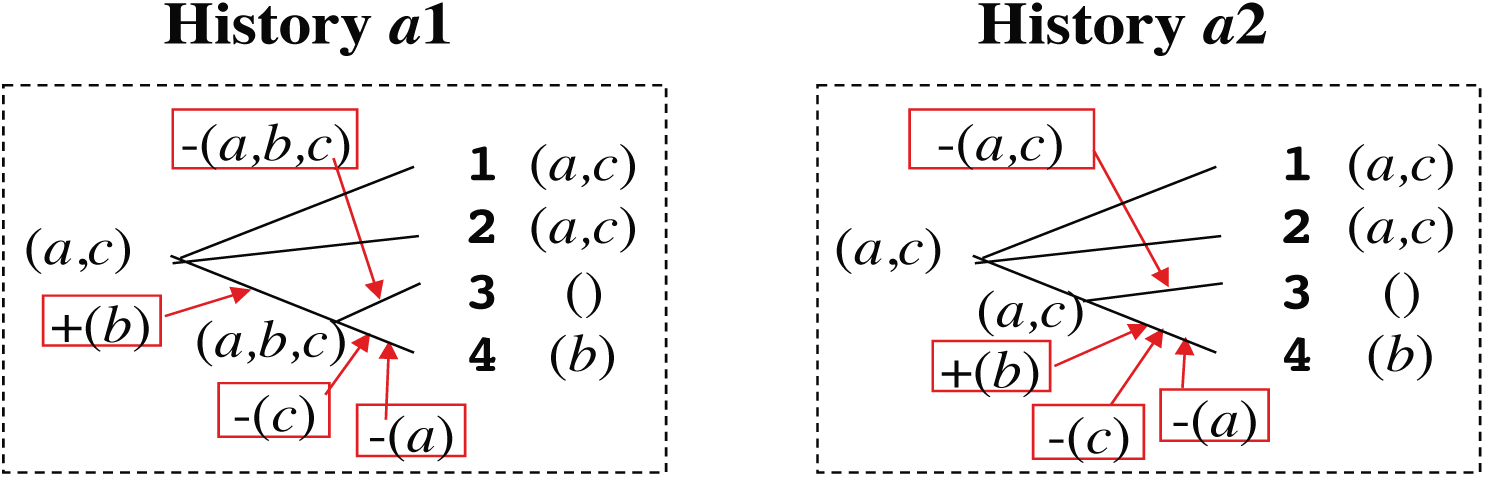
Local indel histories that are likely to actually result in gapped segment as in Figure 7 K. See Methods M2.1 for details.

Then, we simulated MSAs using Dawg (Cartwright 2005), which is known to satisfy a number of criteria as a genuine simulator of the (neutral) evolutionary processes via insertions/deletions (Strope et al. 2009). We created three sets of input MSAs. Sets **1A** and **1B** are homogeneous. Each of them consists of 100,000 MSAs simulated along a three-OTU tree of equal branch lengths (short for 1A and medium for 1B). Set **2** is heterogeneous. It consists of 9,900 MSAs each of which consists of 16 sequences. The MSAs in this set were simulated under typical parameter settings in the BAliBASE benchmark MSA database (Thompson et al. 2005; see Figure 28 for the trees and parameters). All simulations were performed under a biologically realistic Zipf power-law distribution of indel lengths. See Methods M2.2 for details on these simulations.

**Figure 28.**
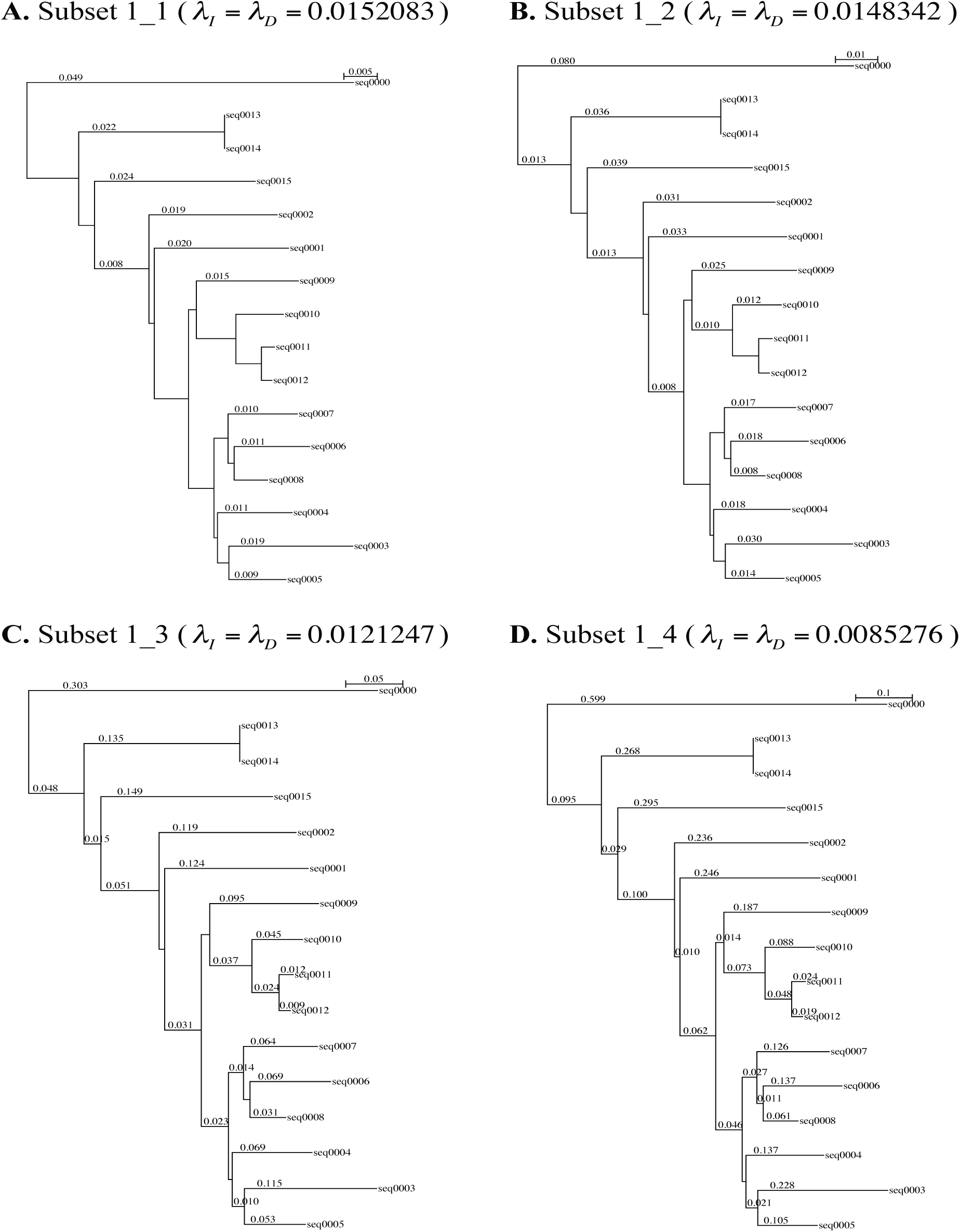
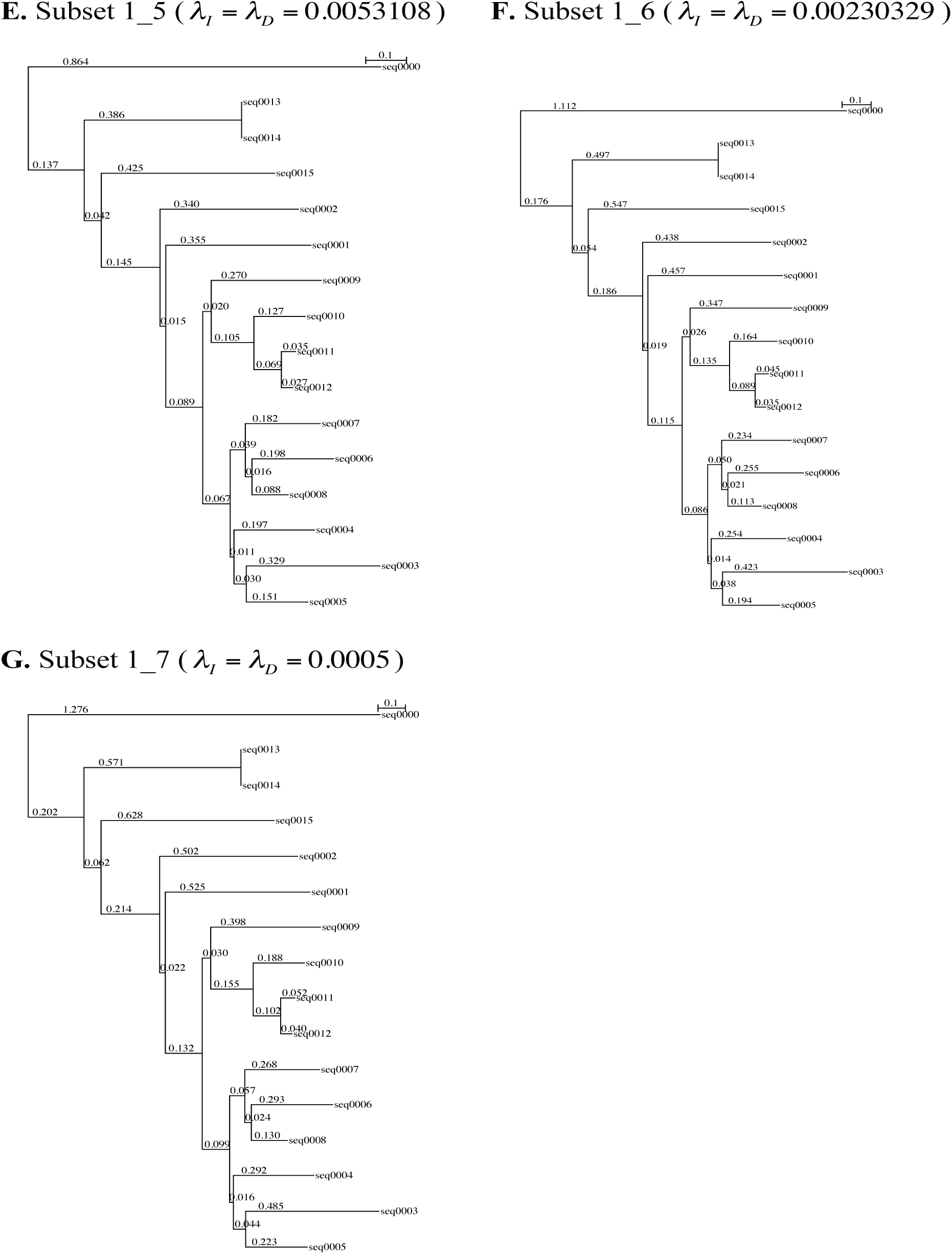
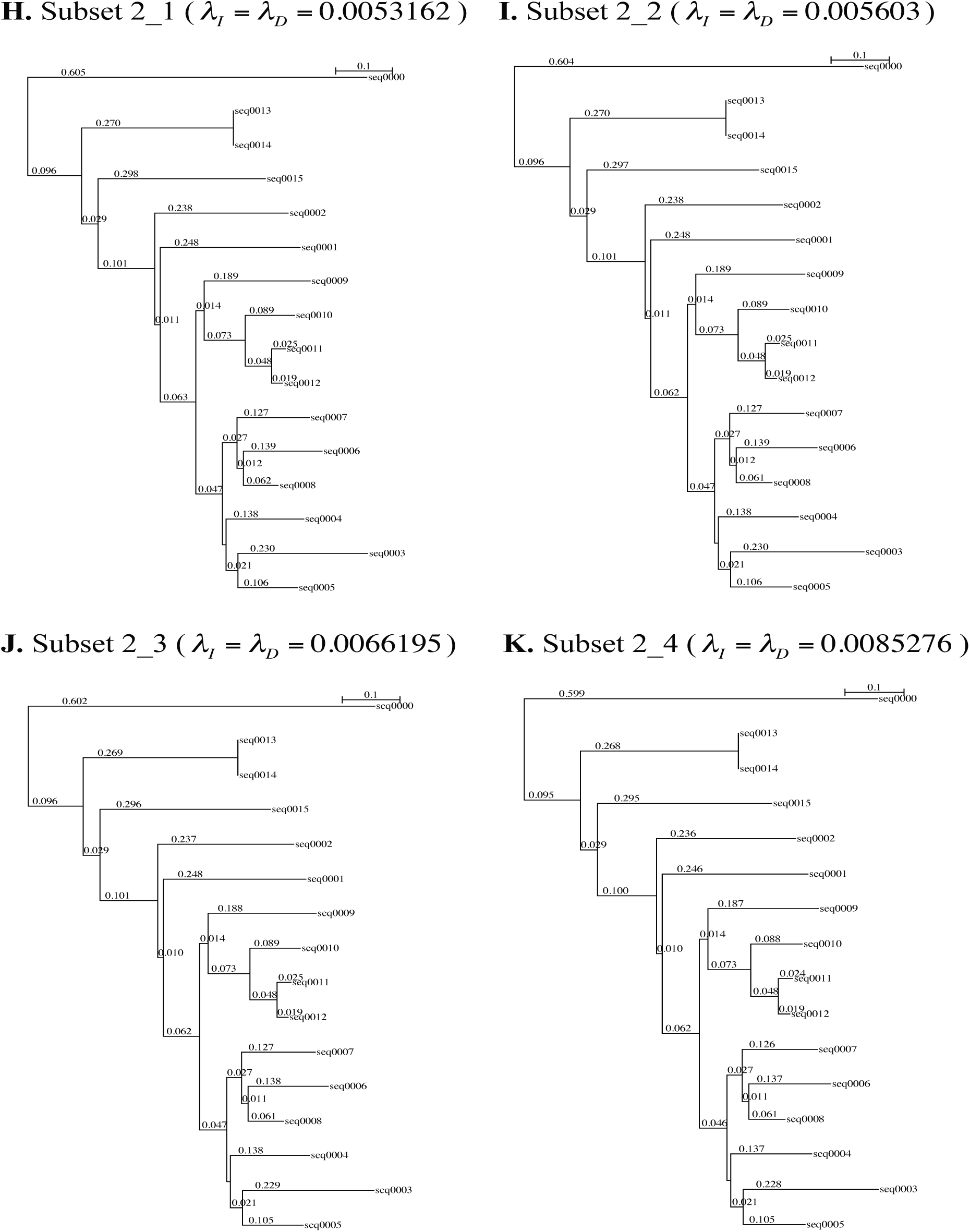
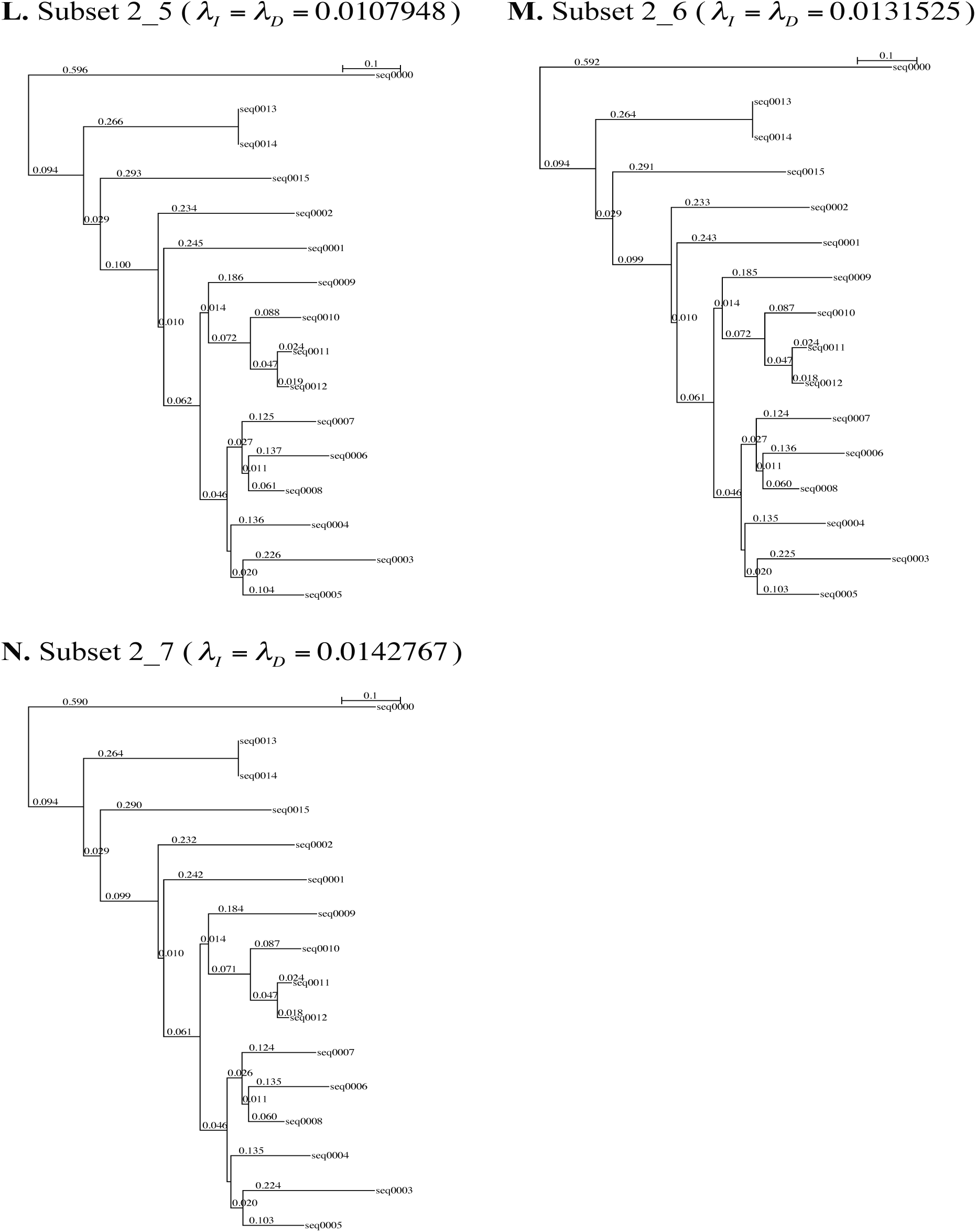
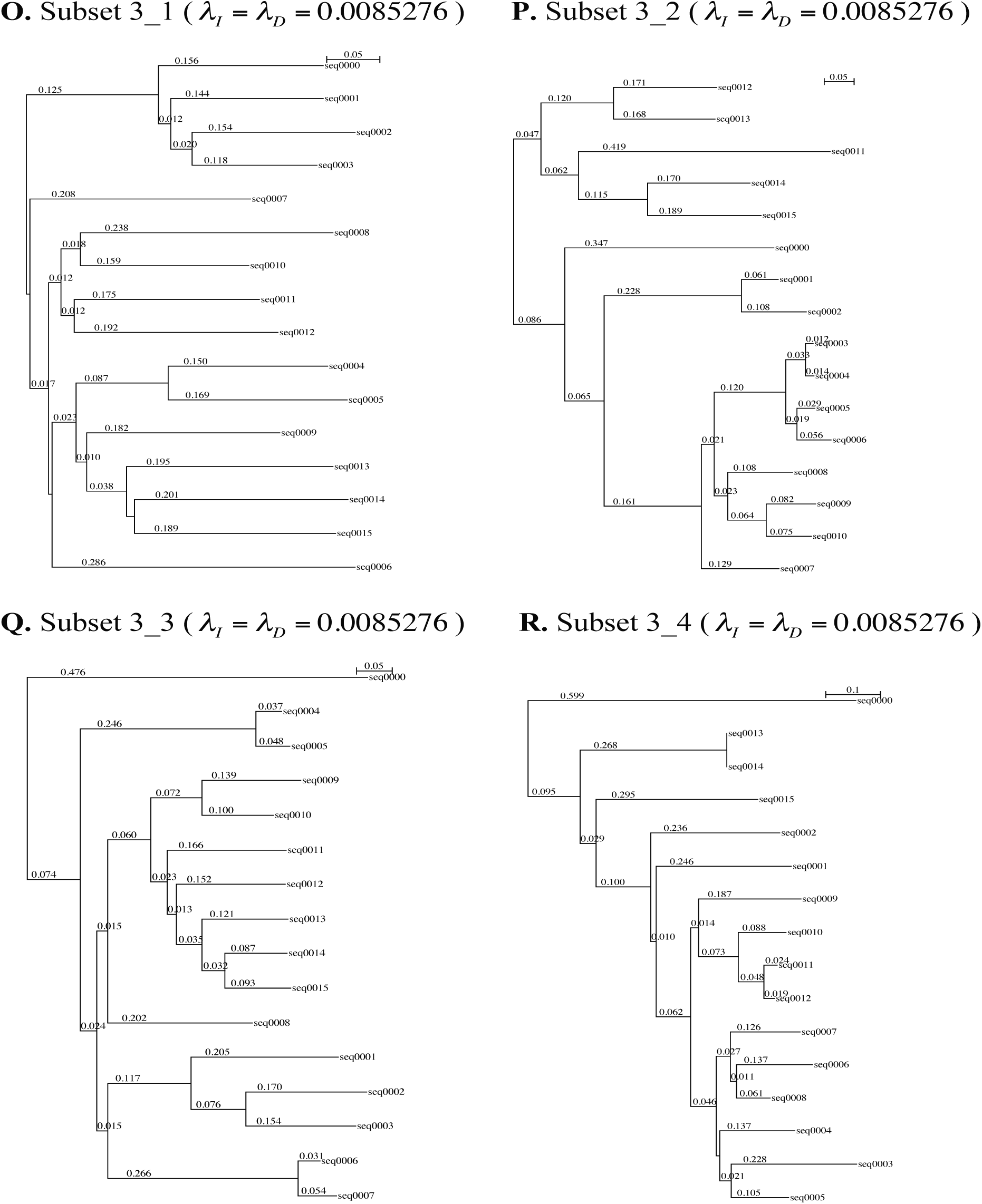
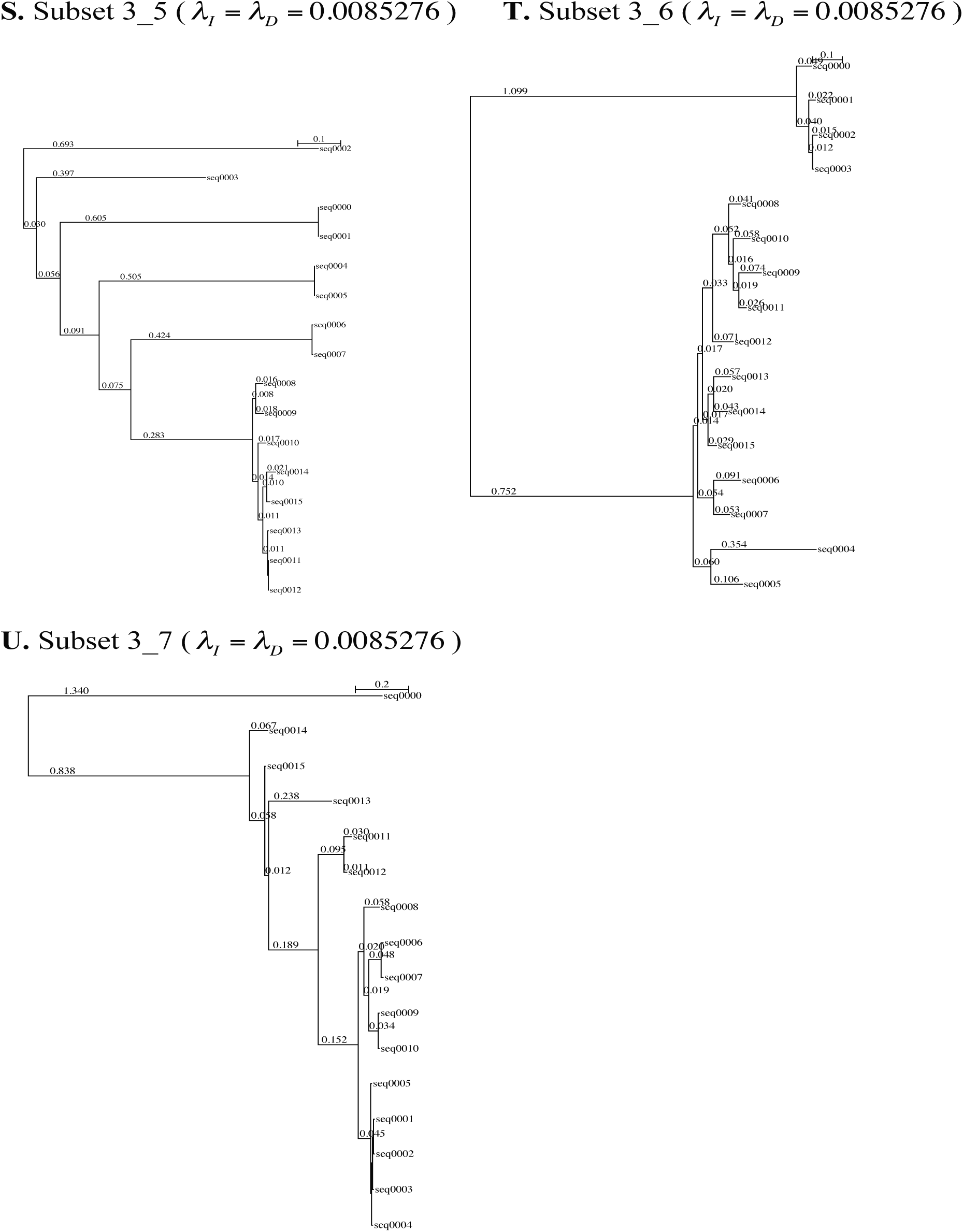
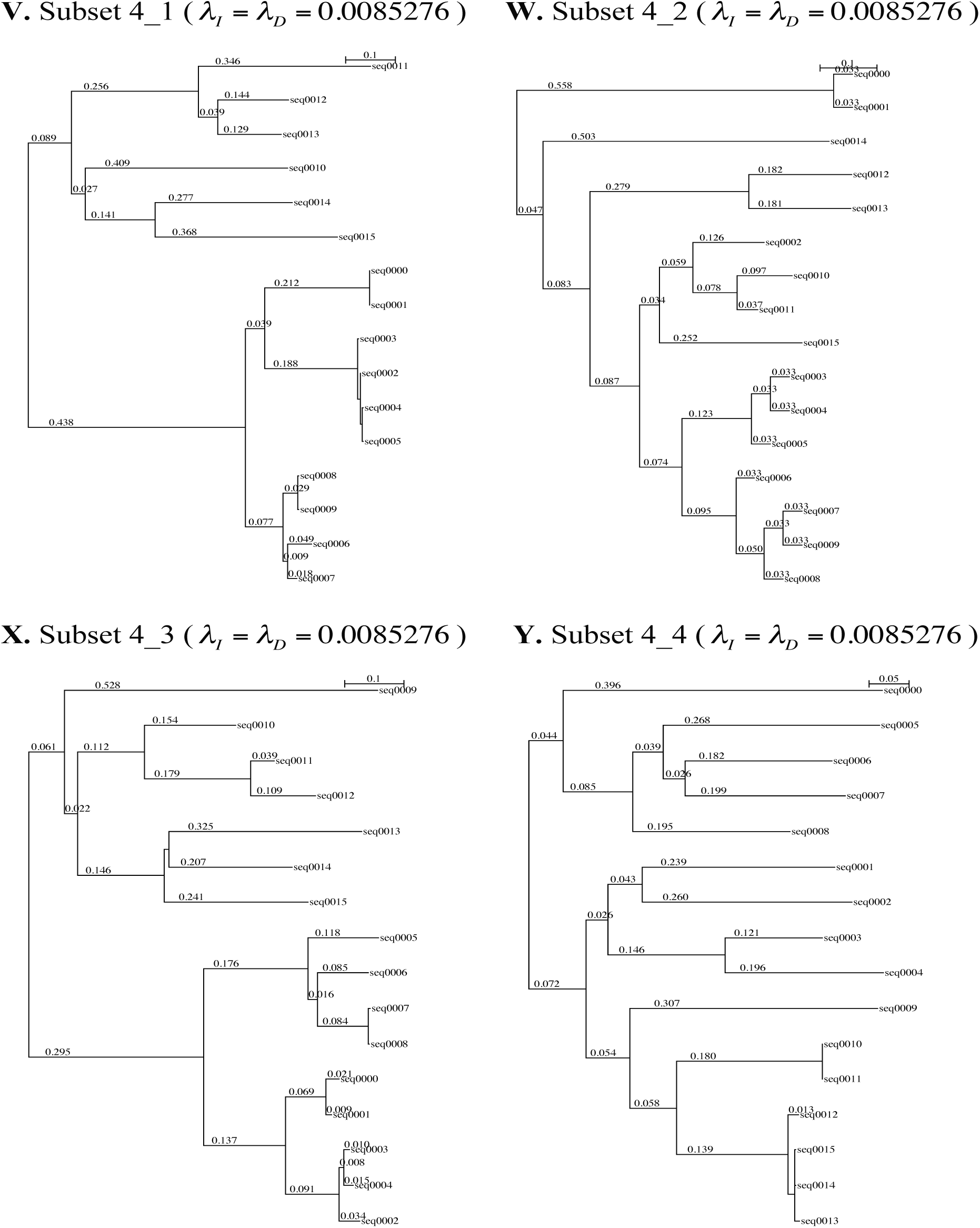
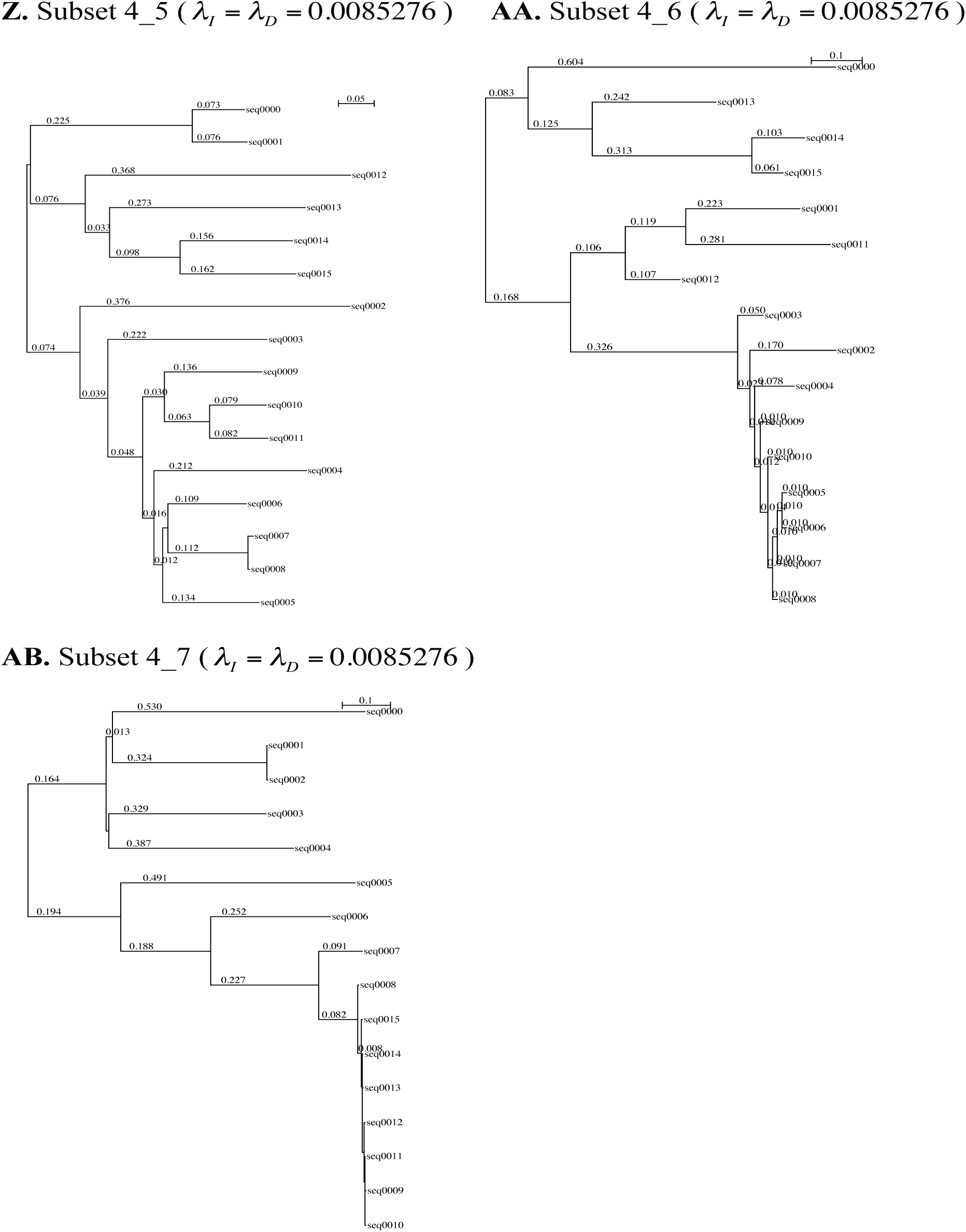
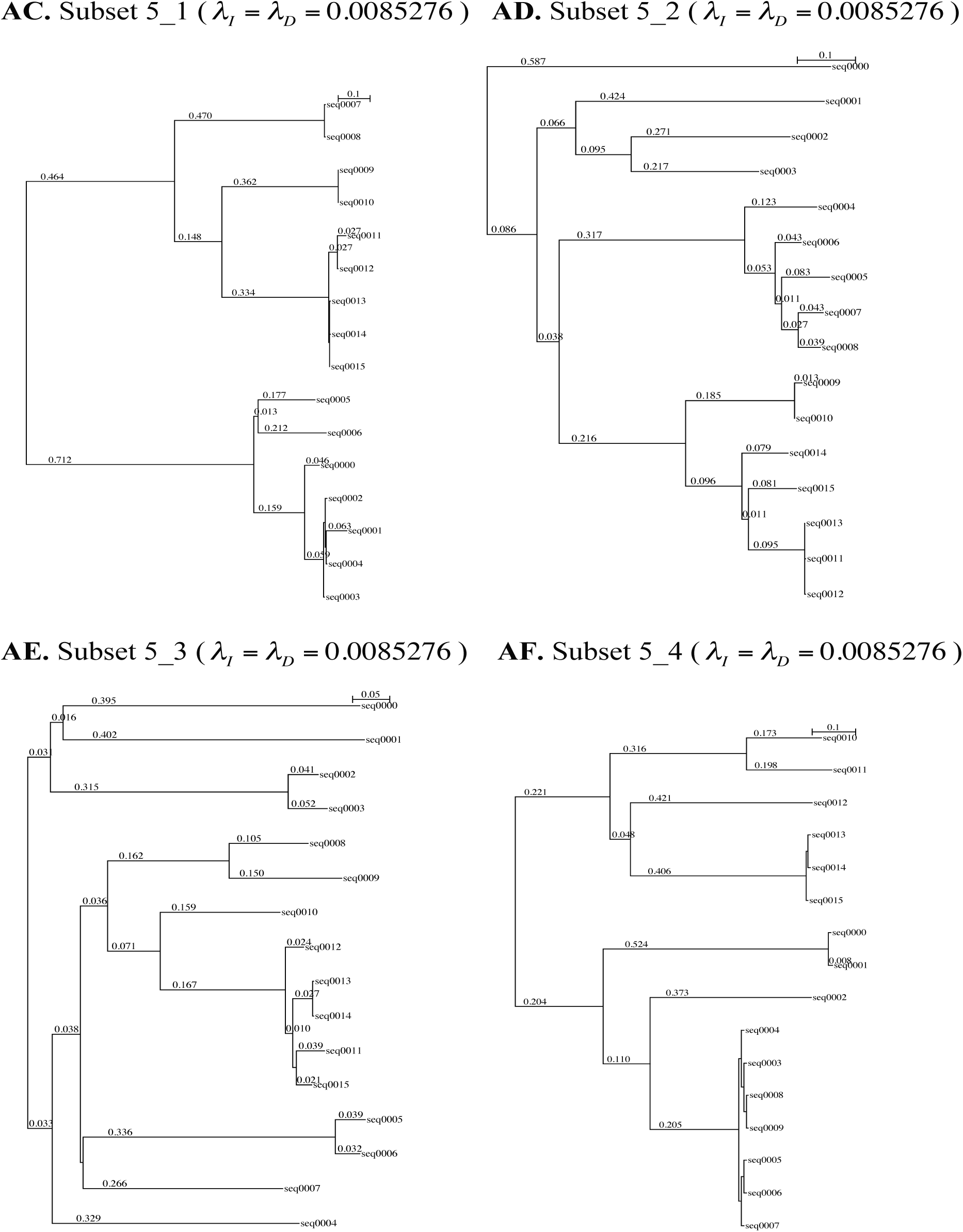
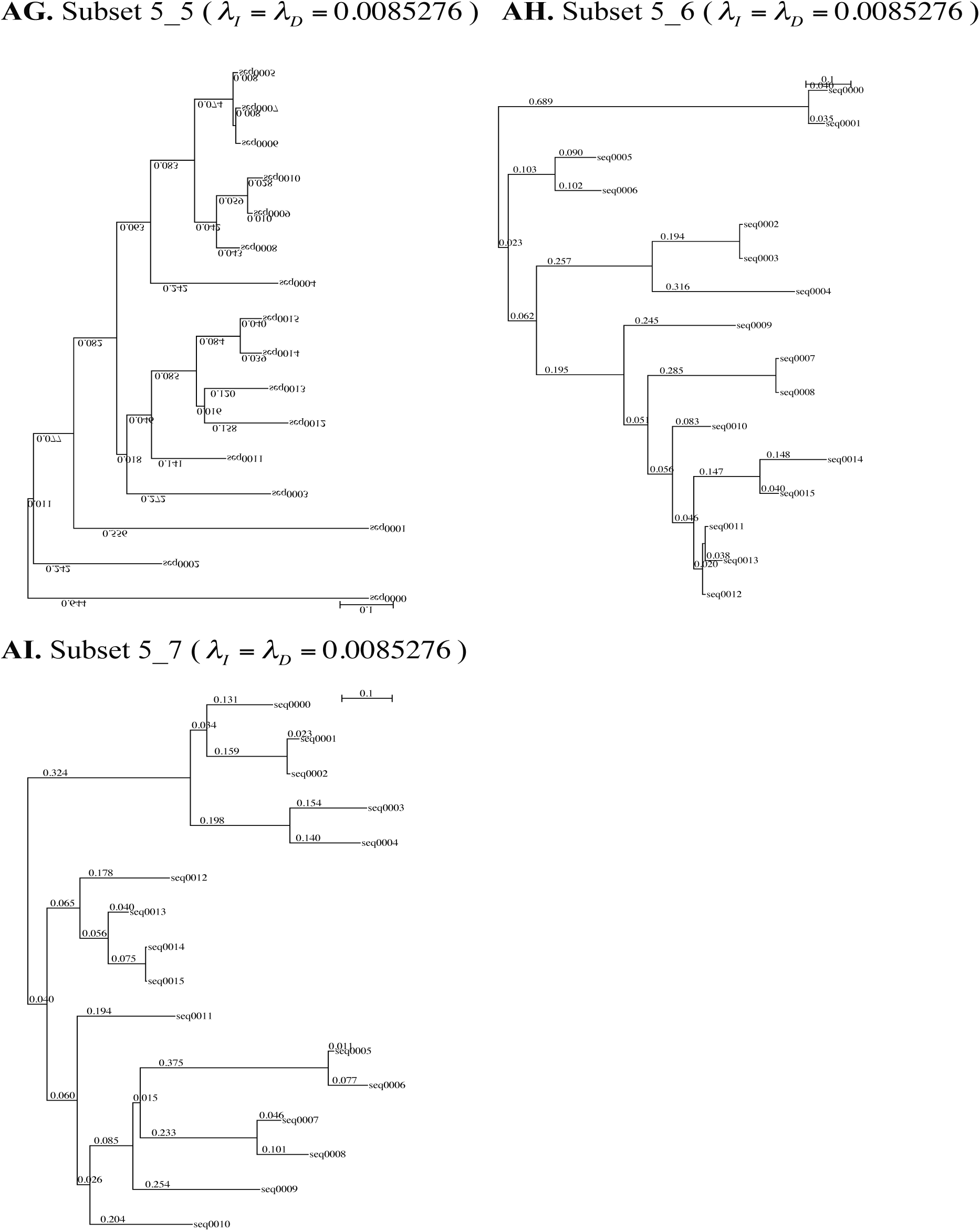
Parameter settings for simulated MSA set 2. Each of the panels A-Z, AA-AI shows the 16-taxon tree and the total rates of insertion and deletion (*λ*_*I*_ and *λ*_*D*_, measured relative to the substitution rate) used for creating a subset of Set 2 of simulated MSAs. Each set is labeled “ *P*_ *L*,” where *P* specifies the parameter varied and *L* specifies the level of the varied parameter. *P* = 1, 2, 3, 4, *and* 5 correspond, respectively, to the mean branch length, the gap content, the coefficient of variation (CV = standard deviation / mean) of the branch lengths, the CV of the number of branches separating a pair of leaves, and the CV of the distance from the root to a leaf. *L* = 1, 2, 3, 4, 5, 6 *and* 7 correspond, respectively, to the 5th, 10th, 25th, 50th, 75th, 90th, and 95th percentiles of the varied parameter among the benchmark MSAs. The remaining 4 out of the 5 parameters were kept at median (*i.e.*, the 50th percentile). Note that the parameters for subsets 2_4 and 3_4 are identical to those for subset 1_4. Thus, to avoid redundancy, we excluded subsets 2_4 and 3_4 from the analysis, and used the remaining 33 subsets for the validation analysis.

The numbers of instances of gapped segments with only gaps of at most 100 bases long were 2676332, 7695575, and 413637 for simulated sets 1A, 1B, and 2, respectively. Out of them, the proportions of instances with non-parsimonious ancestral gap configurations were 0.15%, 1.38%, and 0.33% for the sets 1A, 1B, and 2, respectively. This indicates that non-parsimonious indel histories along the trees contribute only a tiny fraction of the instances of gapped segments. The total number of instances decreased in negative correlations with the length of each gapped segment and the minimum-required number of indels, whereas the relative contribution of non-parsimonious ancestral gap configurations increased in positive correlations with these attributes (data not shown). These results are consistent with the results in Subsection 1.3 of part II (Ezawa, Graur and Landan 2015b).

In each of simulated sets 1A and 1B, we compared the absolute frequency that each local gap-configuration actually occurred with its theoretical prediction, which was based on Eq.(1.1.2a) of part II and included the fewest-indel local histories alone. (See Methods M2.4 for details). Panels A and B of Figure 29 show the scatter plots for the sets 1A and 2B, respectively. The figure indicates that the predicted frequencies with the fewest-indel histories alone (ordinate) are nearly equal to the simulated frequencies (abscissa). Correlation coefficients were 0.9996 for set 1A and 0.9975 for set 1B, and the linear regression analyses yielded relations very close to *Y* = *X* (Table 1). Panel B, however, exhibits a thin downward deviation from the main diagonal around the middle, indicating that the probabilities are underestimated. We confirmed that the deviation disappeared after removing the gap configurations with more than one expected invisible indels (panel C). This is consistent with the results in Subsection 1.3 of part II that the fewest-indel approximation is fairly good as long as the branch lengths and the indel lengths are moderate or shorter.

**Figure 29.**
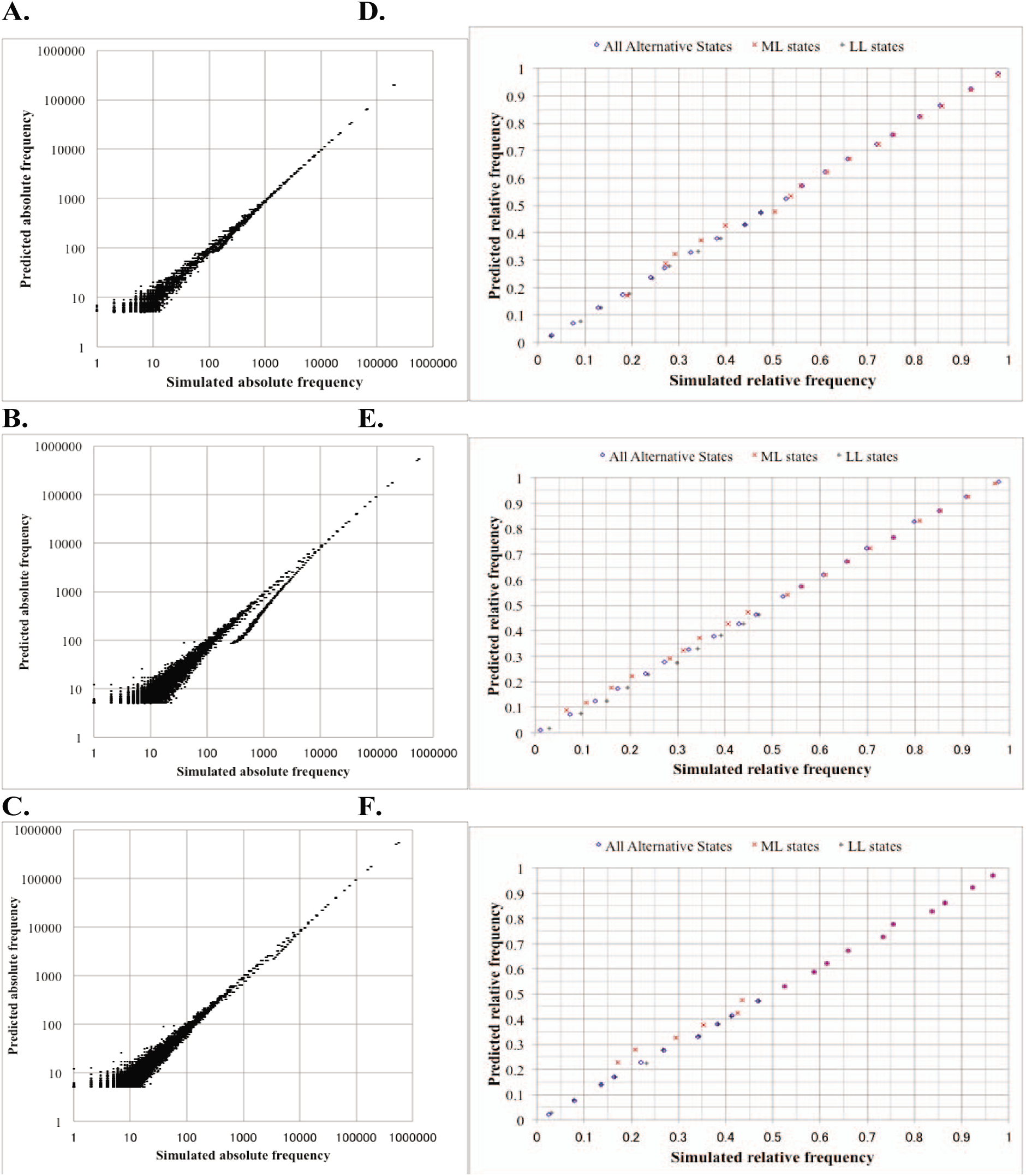
Results of validation analyses via simulations. Each of panels **A**, **B** and **C** compares the predicted absolute frequency of each local gap configuration (ordinate) against the number of times that it actually occurred in a simulated dataset (abscissa). The predicted absolute frequency was calculated using parsimonious local indel histories alone. Note the logarithmic scaling for both axes, which tends to exaggerate sampling errors on the lower-left region in each panel. Panel **A** shows the result with the simulated set 1A. **B.** With set 1B. **C**. With set 1B, after removing long gapped segments. Meanwhile, each of panels **D**, **E** and **F** compares the predicted relative frequencies (ordinate) against the actual relative frequencies in simulations (abscissa). The relative frequencies are among parsimonious local indel histories that potentially yield the same local gap configuration. A blue diamond, a red ‘X,’ and a black cross represent a bin of all parsimonious local indel histories, that of most likely (ML) parsimonious histories, and that of least likely (LL) parsimonious histories, respectively. Panels **D, E and F** show the results with set 1A, with set 1B and with set 2, respectively. See Methods M2.2 through M2.5 for details on the simulation analyses.

**Table 1.**
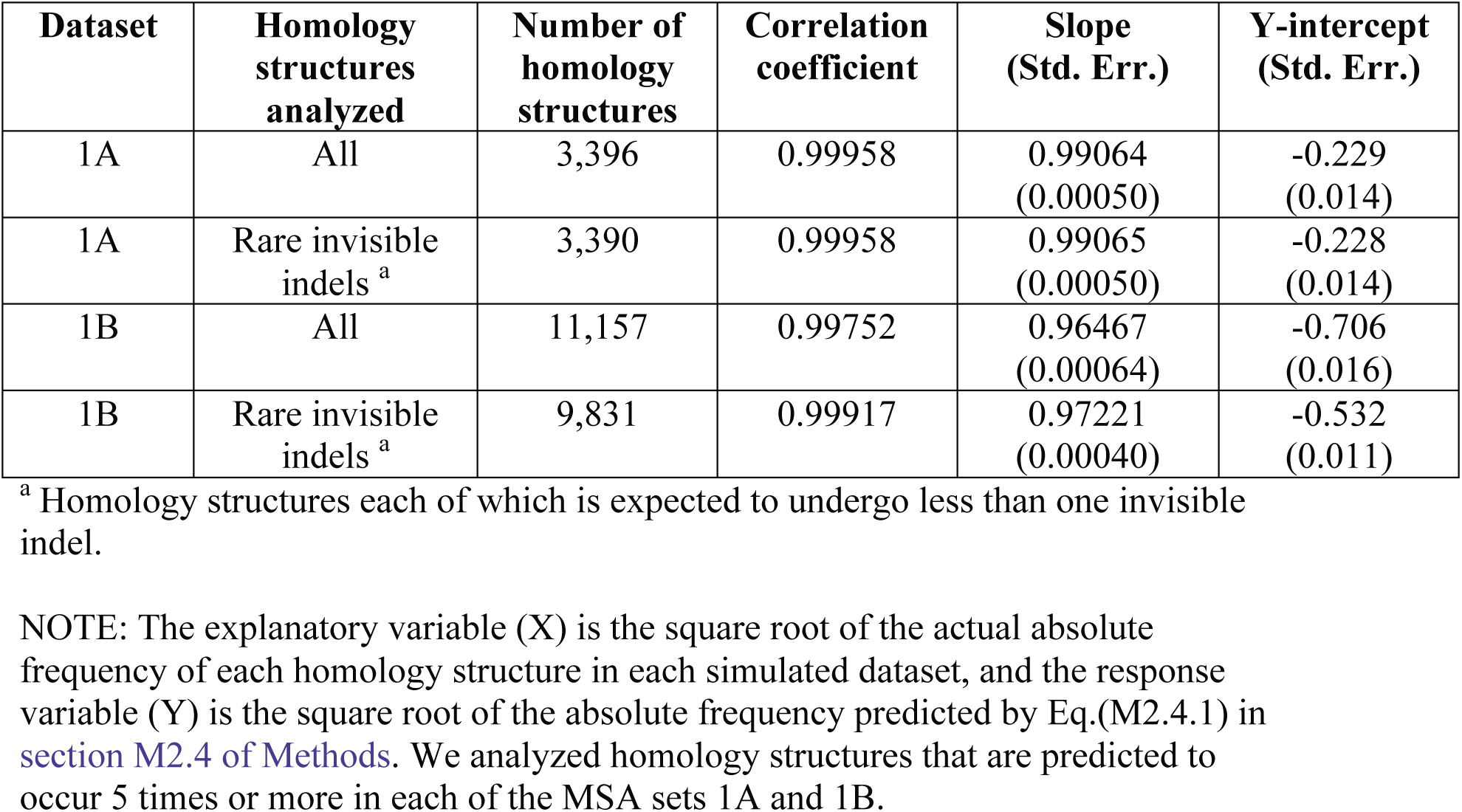
Correlation and regression coefficients between simulated and predicted absolute frequencies of homology structures

| Dataset | Homology structures analyzed | Number of homology structures | Correlation coefficient | Slope (Std. Err.) | Y-intercept (Std. Err.) |
| --- | --- | --- | --- | --- | --- |
| 1A | All | 3,396 | 0.99958 | 0.99064<br>(0.00050) | -0.229<br>(0.014) |
| 1A | Rare invisible indels <sup>a</sup> | 3,390 | 0.99958 | 0.99065<br>(0.00050) | -0.228<br>(0.014) |
| 1B | All | 11,157 | 0.99752 | 0.96467<br>(0.00064) | -0.706<br>(0.016) |
| 1B | Rare invisible indels <sup>a</sup> | 9,831 | 0.99917 | 0.97221<br>(0.00040) | -0.532<br>(0.011) |

<sup>a</sup> Homology structures each of which is expected to undergo less than one invisible indel.
NOTE: The explanatory variable (X) is the square root of the actual absolute frequency of each homology structure in each simulated dataset, and the response variable (Y) is the square root of the absolute frequency predicted by Eq.(M2.4.1) in section M2.4 of Methods. We analyzed homology structures that are predicted to occur 5 times or more in each of the MSA sets 1A and 1B.

Then, we computed the relative simulated frequencies among different sets of ancestral states consistent with the gap configuration of each local MSA (abscissa), and we compared them with their predicted relative probabilities using the fewest-indel local histories alone (ordinate) (Figure 29, panels D, E, and F for sets 1A, 1B, and 2, respectively). See Methods M2.5 for details. The figure demonstrates that the predicted relative probabilities quite well approximated the simulated proportions, whether they are for all alternative histories (blue diamonds), most likely (ML) parsimonious histories (red X’s), and least likely (LL) parsimonious histories (black crosses). Correlation coefficients ranged from 0.99916 to 0.99997, and the linear regression relations were nearly indistinguishable from *Y* = *X* (Table 2). These results collectively imply that the probabilities of the gap-configurations of gapped segments are in general approximated quite accurately even by the contributions from the fewest-indel local histories alone, as long as the branch and the indels are at most moderately long.

**Table 2.** Correlation and regression coefficients between simulated and predicted relative frequencies of correct parsimonious indel histories

| **Dataset** | **Parsimonious indel histories used** | **Number of histories<sup>a</sup>** | **Correlation coefficient** | **Slope (Std. Err.)** | **Y-intercept (Std. Err.)** |
| --- | --- | --- | --- | --- | --- |
| 1A | All | 317,400 | 0.999948 | 1.0105<br>(0.0025) | -0.0045<br>(0.0011) |
| 1A | Most likely (ML) | 119,676 | 0.999793 | 0.9987<br>(0.0049) | 0.0058<br>(0.0038) |
| 1A | Least likely (LL) | 119,856 | 0.999683 | 1.0105<br>(0.0090) | -0.0097<br>(0.0024) |
| 1B | All | 7,252,601 | 0.999967 | 1.0132<br>(0.0019) | -0.00153<br>(0.00023) |
| 1B | Most likely (ML) | 917,499 | 0.999848 | 0.99911<br>(0.00411) | 0.0125<br>(0.0032) |
| 1B | Least likely (LL) | 925,036 | 0.999441 | 0.99905<br>(0.0118) | -0.0136<br>(0.0017) |
| 2 | All | 202,120 | 0.999964 | 1.00083<br>(0.00200) | -0.00089<br>(0.00105) |
| 2 | Most likely (ML) | 91,440 | 0.999156 | 0.9892<br>(0.0096) | 0.0093<br>(0.0084) |
| 2 | Least likely (LL) | 91,440 | 0.999694 | 1.0035<br>(0.0088) | -0.00074<br>(0.00140) |

<sup>a</sup> The number of instances of alternative parsimonious histories of the specified type (All/ML/LL).
NOTE: Here we analyzed all instances of gapped segments in each of which one of 2 or more parsimonious indel histories was “correct.” The explanatory variable (X) is the simulated proportion that the alternative parsimonious indel histories, in each of the 20 bins, actually resulted in the corresponding gap-configurations (“simulated relative frequency”). The response variable (Y) is the average of the predicted relative probabilities of the histories in each bin (“predicted relative frequency”). Note that weighted analyses were conducted. For details, see Section M2.5 of Methods.

Incidentally, for the analysis on relative probabilities, we only used gapped segments each of which can result from two or more parsimonious local indel histories. The instances of such gapped segments accounted for 4.5%, 12.0%, and 22.1% of all “parsimonious” instances in set 1A, set1B, and set 2, respectively. Out of them, the most likely (ML) histories were wrong in 42.3%, 43.4%, and 21.8% of the instances in sets 1A, 1B, and 2, respectively. Therefore, even if we assume that the aforementioned “non-parsimonious” instances were all due to the ML histories that are non-parsimonious, any algorithm that searches for a single ML history would have overlooked the true indel history in 1.9%, 5.2%, and 4.8% of the cases in sets 1A, 1B, and 2, respectively. These frequencies are much larger than those of the “non-parsimonious” instances, *i.e.,* 0.15%, 1.4%, and 0.33% in sets 1A, 1B, and 2, respectively. This indicates that, given correct MSAs and correct trees, our algorithm can recover the true indel histories more frequently than any algorithm to search for a *single* ML history.

## Discussion

In the following, we will discuss possible improvements in our algorithms (D1), risks associated with the naïve applications of our algorithm (D2), and possible applications of our theory and algorithms (D3).

### D1. Improvements in our algorithms

In order to show that the lowest-order (*i.e*., fewest-indel or parsimonious) terms of perturbation expansion approximate the alignment probability fairly well, we developed an algorithm that enumerates all fewest-indel histories and calculates their contributions to the alignment probability. However, the current version of the algorithm is still rudimentary and there are some rooms for improvements. Some mandatory improvements would be the incorporation of regional and lineage-wise variations in the indel rate parameters, the incorporation of indel length distributions other than the geometric and power-law distributions, and the implementation of a function to estimate indel model parameters from input data. These features will be coming soon. Moreover, the following improvements would be worth pursuing.

#### D1.1. Enabling to handle long gaps

In the simulation analyses in this paper, we only analyzed gapped segments that are at most 100 bases long. This is because longer gapped segments could result from millions of fewest-indel local histories, and thus it could take too long for our current version of the algorithm to finish the analysis of millions of gapped segments that are necessary for assessing the accuracy of the estimated probabilities. Here let us briefly estimate the time complexity of the component of our current algorithm to handle each gapped segment. The component consists of sub-components (for details, see Methods M1 and Figure 1). (i) It first partitions the segment into blocks of contiguous columns with the same gap pattern. (ii-a) It second constructs an initial candidate of the fewest-indel local history. This is done by identifying the Dollo parsimonious indel history for each block, and by concatenating insertions or deletions along a same branch and in effectively contiguous blocks of distinct gap patterns. (ii-b) Then, starting with the initial candidate, it attempts to enumerate all possible fewest-indel histories, by trying the “branch-and-merge” operation on every indel event in each candidate history. And finally, (iii) it calculates the total occurrence probability of the gapped segment, by calculating the probability of each local history and by summing the probabilities over all fewest-indel local histories. The time complexity of (i) is roughly *O* (*N*^*X*^ *L*(*C*_K_)), where *N*^*X*^ is the number of sequences in the MSA and *L*(*C*_K_) is the number of columns in the gapped segment (*C*_K_). Sub-component (ii-a) has a rough time complexity *O*(*N*^*X*^ *B*), where *B* is the number of blocks in the segment. Sub-component (ii-b) ends as soon as the algorithm can find no further candidate with indels fewer than or as few as those in each current candidate. Thus, the time complexity of (ii-b) is roughly bounded from above by 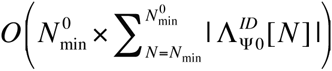. Here 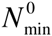 is the number of indels in the initial candidate history, and 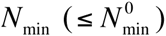 is the minimum number of indels necessary for the gapped segment. And 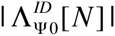 denotes the number of local histories, each of which results in the segment via *N* indels, with the minimum number of indels along each branch and with no “null indel histories.” Finally, the time complexity of (iii) is roughly at most 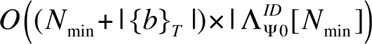, where |{*b*}_*T*_| is the number of branches. From these arguments, we realize that sub-components (i) and (ii-a) can be performed faster than the common algorithms to calculate the likelihood of the residue configuration under an independent-site substitution model (*e.g*., Felsenstein 1981, 2004; Yang 2006). We also realize that 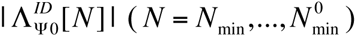 hold the key to the time-complexity of sub-components (ii-b) and (iii), and thus of the entire algorithm. Because *N*_min_ is in general quite close to 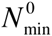, and because 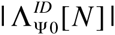 is expected to increase with *N*, it would be enough to estimate 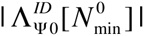. A typical situation with an extremely large 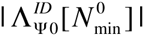 is when a segment contains a long gap and many short gaps. Here, it should be noted that each “branch-and-merge” operation works on a pair of indels along neighboring branches. Thus, even if the segment includes multiple indels that are spatially overlapping each other, as long as they are along branches not neighboring each other, they are usually not subject to the “branch-and-merge” operation. Taken together, these arguments suggest that the most important situations are where there is a long insertion along a branch whose child branches hold many short gaps, and, alternatively, where there is a long deletion along a branch whose parent and/or sibling branches hold many short gaps. Because each short gap is explained by two alternative fewest-indel histories (a deletion along one branch and an insertion along the other), if there are *NShG* such short gaps, 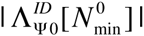 is roughly *O*(2^*N*_*ShG*_^). And 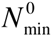 should in general be less than *N*^*X*^ *B* and greater than *B*/2. Thus, we expect that the time complexity will be typically from *O*(*B*· 2^*N*_*ShG*_^) to *O*((*N*^*X*^ *B*)^2^ · 2^*N*_*ShG*_^). For example, when *B* = *N*_*ShG*_ = 20, the time complexity is larger than *O*(20·2^20^) ≈ *O*(2 · 10^7^), and the algorithm is expected to consume a large amount of time. The expected value of *N*_*ShG*_ is roughly (*λ*_*I*_ + *λ*_*D*_)(|*b*_1_| + |*b*_2_|) *L*(*C*_K_), where (*λ*_*I*_ + *λ*_*D*_) is the total rate of indels per substitutions, and | *b*_1_ | and | *b*_2_ | are the lengths of the child branches (in the unit of the expected number of substitutions per site) in the case of a long insertion along a branch. For example, if *λ*_*I*_ + *λ*_*D*_ takes an upper-bound of 0.2, and when *L*(*C*_K_) = 200, *N*_*ShG*_ is expected to be around 40 ×(| *b*_1_ | + | *b*_2_ |), which is 20 if | *b*_1_ | + | *b*_2_ | = 0.5. Thus far, we only considered a case with a simple gap configuration. However, the essence of the problem should remain unchanged even when we deal with a gapped segment with a more complex gap configuration and/or with many sequences. In such a case, the time complexity depends heavily on the total length of the long gaps and the lengths of the neighboring branches, but not directly on the number of sequences. Rather, if evenly sampled along the tree, an increased number of sequences could reduce the time complexity, thanks to the decreased branch lengths. This reasoning implies that one way to reduce the expected value of *N*_*ShG*_ would be to densely sample the sequences so that the branch lengths will decrease. Such a sample, however, will not always be available.

Thus, in order to handle a long gapped segment in a reasonable amount of time, some technique needs to be devised or borrowed from somewhere. One promising technique would be to hierarchically handle gaps, first long ones only and then short to medium ones. More precisely, by chopping a MSA into a number of sub-segments according to the configuration of long gaps alone (panel B of Figure 30), we can perform the original task in two steps. (1) We first infer the broad histories consisting of long indels alone and calculate their approximate probabilities (panel C). And (2) after ignoring branches of long gaps, we further infer the “fine-grained” histories of short to medium indels in each sub-segment and calculate their approximate probabilities (panel D). For example, let us consider that the method is applied to the aforementioned example case of a long indel with *N*_*ShG*_ = 20 overlapping short indels along neighboring branches. Then, the time complexity would be reduced from greater than *O*(10^7^) to approximately *O*(20 × 2) < *O*(10^2^). A similar sub-division strategy was proven to work fairly well on the algorithm that singles out a parsimonious indel history when one or more long gaps are involved (Chindelevitch et al. 2006). Thus, we hope that the strategy proposed here will work at least reasonably also to the algorithm presented in this paper.

**Figure 30.**
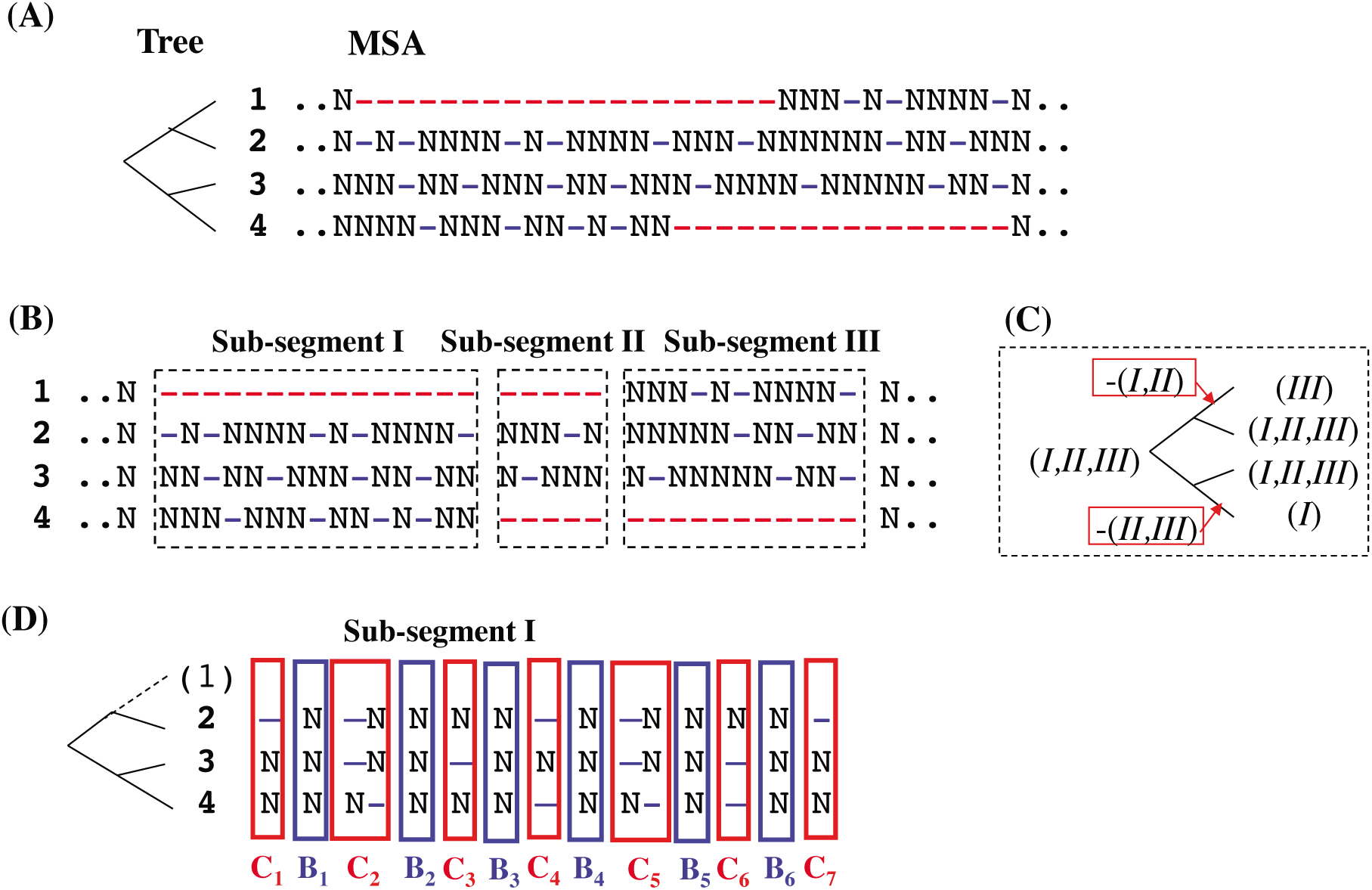
Problem with long gaps and its solution by hierarchical partitioning. **A.** A gapped segment with long gaps (red) often contains lots of short gaps (blue), which makes the simple partitioning less effective. **B.** A coarse-grained partitioning according only to the configuration of long gaps, chopping a gapped segment into a number of sub-segments (I, II and III in this example). **C.** A broad indel history resulting in the configuration of long gaps. **D.** Fine-grained partitioning of the Sub-segment I in panel B according to the configurations of short gaps. This measure decomposes the problem with potentially numerous indel histories that need be considered into a set of sub-problems, each of which has at most a few candidate histories. Ignoring the sequence containing the long gap enables this measure. The ignored sequence is indicated by the parenthesized sequence ID and the dotted branch leading to it. Each column in this figure should be regarded as a gap-pattern block rather than a single site.

Another problem is that, as a gapped segment gets longer, its probability gets less accurately approximated by the sum of probabilities of the fewest-indel local histories alone, as shown in Subsections 1.2 and 1.3 of part II (Ezawa, Graur and Landan 2015b) as well as in Results of this paper. The solution to this problem will be discussed in the next subsection.

#### D1.2. Incorporating local histories with more than fewest indels

As shown in Subsections 1.2 and 1.3 of part II and in Results of this paper, the fewest-indel local indel histories predominate only when the gapped segment and branches in the tree are at most moderately long. Thus, the approximation by these histories alone gets less and less accurate as the segment and/or the branches get longer and longer. Moreover, in terms of the *absolute* frequency, a next-fewest-indel local history with two indels should occur more frequently than a fewest-indel local history with, *e.g*., four indels. If such cases need to be included into the analysis, it would become inevitable to incorporate local histories with more than the fewest indels, *i.e.*, sub-parsimonious local indel histories.

According to the local MSA version of Eqs.(3.2.13a,b’) of part I (Ezawa, Graur and Landan 2015a), elements of sub-parsimonious local indel histories can be classified into two broad types. One consists of (A) sub-parsimonious histories along a branch, yielding higher-order corrections to the PWA probability along the branch. And the other consists of (B) sub-parsimonious histories yielding sets of basic states at interior nodes that are different from any of those for the parsimonious histories. In Subsection 1.2 of part II (Ezawa, Graur and Landan 2015b), the histories of broad type (A) were examined quite thoroughly, up to the next-fewest indel histories for all gap configurations, and up to “all orders of perturbation” for all but one type of gap configurations. Thus, if desired, it is possible to incorporate this type of histories. Although their calculation will take longer than the calculation of the fewest-indel terms, we could use pre-computed multiplication factors, as long as we know the branch lengths and indel model parameters. By using such pre-computed multiplication factors for the PWA probabilities that take account of higher order terms, the accuracy of the local MSA probabilities will considerably improve even if we ignore indel histories of broad type (B), as indicated in Subsection 1.3 of part II. Taking account of broad type (B) would require an algorithm to systematically enumerate such local indel histories. Some hints on such an algorithm would come from the “branch-and-merge” operation (Methods M1.2), and from the examples considered in Subsection 1.3 of part II and Appendix A2 of part II. As we see, they exemplify how type-(B) next-fewest-indel histories could be constructed from a fewest-indel history.

Incidentally, it would be worth a mention here that our parsimony algorithm is not yet perfect, in the sense that the current version cannot find some deletion-dominated local histories creating gap-configurations belonging to the class of “intersection between cousins” described by Fredslund et al. (2003). (For more details, see Methods M2.1.) Nevertheless, this drawback will only slightly, if at all, impact the overall performance of our algorithm, because such local histories should be very rare, with the probabilities at most on the order of (*r*_*D*_ × | *b* |)^4^, which is typically less than 10^−7^. However, if we are interested, *e.g*., in indel hot spots where multiple indels stack together, this drawback may lead to erroneous pictures. We hope to incorporate the function to find such histories in the near future, maybe along with the extension to handle long gapped segments.

### D2. Risks associated with naïve applications of our algorithm to *reconstructed* alignments

Some readers might have thought that they would be able to predict the (local) indel histories, as well as the accompanying sets of ancestral gap configurations, quite accurately by applying the algorithm presented here to a MSA *reconstructed* by one of the state-of-the-art aligners (reviewed, *e.g*., in Notredame 2007). On the contrary, we caution the readers that such analyses would be premature. What we demonstrated in this paper is that our algorithm gives quite accurate estimations of the occurrence probabilities of gapped segments, as well as of the relative probabilities among fewest-indel histories giving rise to each gapped segment, *provided that* it is fed a *correct* MSA. Unfortunately, however, recent analyses (*e.g*., Löytynoja and Goldman 2008; Landan and Graur 2009) showed that reconstructed MSAs are considerably error-prone, *even if they were reconstructed via state-of-the-art aligners.* This means that a naïve application of the current version of our algorithm to a *reconstructed* MSA is fraught with high risks of incorrect predictions of indel histories, because MSA errors immediately lead to erroneous reconstructions of ancestral gap configurations. We therefore advise readers to avoid such a kind of analyses whenever possible. Or, even if they do, the possibilities of MSA errors must be fully taken into account when interpreting the results. Originally, the theoretical formulation and the algorithm presented in this paper were developed for the purpose of comparing candidate MSAs in terms of their occurrence probabilities. This will be among the subjects of the next subsection.

### D3. Possible applications of our theory and algorithm

As suggested by the results in this paper, our theoretical formulation is applicable to examining the parameter regions within which existing indel models are likely to work. And the formulation was also shown to suggest possible modifications and extensions of the models so that they will better approximate *genuine* and biologically realistic evolutionary models. In addition to them, here we will discuss some other areas that it and the algorithm presented in this paper may possibly be applicable to.

First, as originally intended, our methods will be used to identify the most likely MSA among multiple candidate MSAs. Or, if coupled with a smart MSA sampler that can preferentially explore regions of the MSA space where the MSAs are quite likely, our methods could provide a “probability distribution” of quite likely MSAs. The question is whether or not such a MSA sampler is available or, if still unavailable, can be constructed. In Appendix A1 of part I (Ezawa, Graur and Landan 2015a), we demonstrated that the method of Miklós et al. (2004) is equivalent to our *ab initio* method when calculating the probability of each LHS equivalence class of indel histories during a time-interval. This equivalence holds certainly under space-homogeneous indel models and possibly under the model provided in Subsection 5.3 of part I. This implies that, at least under these models, the dynamic-programming (DP) algorithm provided by Miklós et al. (2004) could be applicable also to our *ab initio* calculation of the PWA probabilities. The problem is that their DP algorithm might be too slow to be applicable to the sampling of MSA probabilities. Some recent probabilistic MSA aligners (*e.g*., Paten et al. 2008; Westesson et al. 2012) provides an approximate probability distribution of MSAs with a reasonable time complexity by exploring only a neighborhood of a MSA reconstructed by an optimum-search-type aligner. It is an open question whether or not a similar MSA space exploration strategy, or any other smart exploration strategies, can be coupled to our algorithm. If this is the case, our algorithm could potentially predict a MSA probability distribution more accurately than the previous ones. This is because ours can accommodate biologically realistic features, such as power-law indel length distributions, whereas the previous ones are based on (nearly) standard HMMs or transducers and thus can only accommodate geometric distributions. Past studies on the stochastic pairwise sequence analyses revealed that taking account of realistic indel length distributions improves the accuracies and consistencies of the analyses (Lunter et al. 2008; Cartwright 2009). And we expect that this will also be the case with the estimation of a MSA distribution. Once we obtain a fairly accurate MSA probability distribution, then, we will be able to reconstruct (the probability distribution of) the ancestral gap configurations quite accurately, as desired by many researchers.

Thus far, we only discussed the applications of our theoretical formulation and our algorithm to the situations where a correct phylogenetic tree is known. In real sequence data analyses, however, as opposed to those based on simulations, we do not know the correct tree in advance, and thus a phylogenetic tree must also be inferred from the sequence data. A commonly employed practice to infer a phylogenetic tree is as follows: first a rough guide tree is constructed by applying a fast phylogeny reconstruction method to the result of pairwise sequence comparisons; then an MSA is reconstructed with the aid of the guide tree; and finally a purportedly more accurate phylogenetic tree is inferred using the reconstructed MSA as an input. However, phylogenetic trees inferred in this way tend to be incorrect and biased toward the rough guide trees initially constructed (see, *e.g*., the introductions of Lunter et al. (2005) and Redelings and Suchard (2005)). Moreover, as mentioned in Discussion D2, a single reconstructed MSA is also error-prone. Thus, at least theoretically, an ideal way would be to simultaneously estimate the MSA and the phylogenetic tree from a set of homologous sequences. Or, an even better way would be to infer a joint distribution of MSAs and phylogenetic trees. To the best of our knowledge, Holmes and Bruno (2001) first developed such an algorithm. Later, several measures were devised to accelerate the analysis (*e.g*., Lunter et al. 2005; Redelings and Suchard 2005, 2007; Suchard and Redelings 2006; Novak et al. 2008). Still, it takes a considerable amount of time to perform an analysis (but see also Bouchard-Côté and Jordan 2013). It would be desirable if our algorithm can be applied to the simultaneous estimation of the MSA and the tree with a reasonable time complexity. This would undoubtedly be a formidable problem, given the large time consumption of the previous algorithms cited above, despite the fact that they are based on single-residue indel models or geometric indel length distributions and thus are undoubtedly more efficient than our current algorithm. Although the implementation of the measure discussed in Discussion D1.1 would substantially reduce the consumed time, this alone would not be enough to realize a practical tool. A key to the success would be whether or not the measures devised in the previous studies, or their modified versions, could be combined with our algorithm.

Another possible application of our algorithm would be to the inference of a phylogenetic tree when a MSA is input. Because shared indels are phylogenetically informative, they could in principle be used to reconstruct a phylogenetic tree (*e.g*., Rivas and Eddy 2008). Particularly, they could be useful in a situation where indels occur nearly as frequently as substitutions and, at the same time, where branches are too short to accumulate enough substitutions to resolve the phylogenetic relationships (*e.g*., Redelings and Suchard 2007). Extra caution should be advised, however, when using indels to infer phylogeny from a *reconstructed* MSA, because it is highly prone to errors. Such errors could immediately mislead the prediction of shared indels (see Discussion D2). It may nevertheless be somewhat useful to develop an algorithm to incorporate indel information to the inference of a sequence phylogeny. Traditionally, the maximum likelihood (ML) inference of the phylogenetic tree was based on residue configurations of the columns in a MSA under a given substitution model. For this purpose, some useful heuristic tree search methods, such as nearest-neighbor interchanges (NNI) and sub-tree pruning and re-grafting (SPR), have been devised (see, *e.g*., Felsenstein 1981, 2004; Yang 2006). It may be worth trying to adapt our algorithm to the techniques developed thus far for the inference of the substitution-based ML phylogeny (*ibid*.), and also to the methods used in the previous attempts to use indels for the ML phylogenetic tree reconstruction (*e.g*., Rivas and Eddy 2008). A key point would be whether or not our algorithm, especially with the feature discussed in Discussion D1.1, can be modified to enable an efficient computation of the probabilities of gapped segments after each NNI or SPR step. We invite the interested readers to tackle the problems in this section, as these problems are completely open to future studies.

## Conclusions

In a previous study (Ezawa, Graur, and Landan 2015a), we established the theoretical basis of an *ab initio* perturbative formulation of a general continuous-time Markov model, which is a *genuine* stochastic model describing the evolution of an *entire* sequence via indels along the time axis. Using the formulation, we proved that, under a certain set of conditions, the *ab initio* probability of an alignment is factorable into the product of an overall factor and contributions from local alignments delimited by preserved ancestral sites (PASs). In another previous study (Ezawa, Graur, and Landan 2015b), we showed how we can concretely calculate the fewest-indel contributions, and also the next-fewest-indel contributions, to the probability of each local alignment.

Based on these results, here in this study, we developed an algorithm that calculates the first approximation of the probability of an input multiple sequence alignment (MSA), given a parameter setting including a phylogenetic tree. The algorithm does this job by summing the contributions from *all* fewest-indel histories (*i.e*., parsimonious indel histories) consistent with the MSA. We performed some validation analyses using a *genuine* molecular evolution simulator, Dawg (Cartwright 2005). The results indicated that even the first approximation can estimate the true probabilities of local MSAs quite accurately, as long as the gaps or tree branches are not too long. This algorithm, along with the analytical methods developed in (Ezawa, Graur, and Landan 2015b), gives us a hope that our *ab initio* perturbative formulation can indeed be practically used, *e.g*., for estimating the reliability of reconstructed MSAs and for reconstructing the ancestral gap configurations given a true MSA. The algorithm and the analytical methods are available as a package named LOLIPOG (<u>lo</u>g-<u>li</u>kelihood for the <u>p</u>attern <u>o</u>f <u>g</u>aps in MSA) at the FTP repository of the Bioinformatics organization (Ezawa 2013).

However, caution should be exercised when directly applying LOLIPOG to the analyses of real biological data, because most of the reconstructed alignments between/among the biological sequences are considerably erroneous, especially regarding the positioning of gaps (*e.g*., Löytynoja and Goldman 2008; Landan and Graur 2009). It would thus be preferable to first develop programs that take advantage of the fruits of this series of study to accurately estimate and rectify the errors of reconstructed alignments under a genuine and biologically realistic evolutionary model of indels.

## Methods

### M1. Algorithm

As briefly mentioned in Results, we developed an algorithm that, under a given phylogenetic tree of the sequences and a given indel model (including its parameters), calculates the first-approximate probability that a given MSA actually occurs, using only the fewest-indel histories (*i.e*., parsimonious indel histories) consistent with the MSA. As a byproduct, the algorithm also calculates the relative probabilities among the parsimonious indel histories. In this section of Methods, we will describe the algorithm. Then, in Section M2, we will describe the analyses that were performed to validate the algorithm.

In this Methods, when we refer to a “MSA,” we will consider only its *gap-configuration* (*i.e*., differences in the residue states will be ignored). For example, the “probability of a MSA” means the probability of *the gap-configuration of* the MSA under a given *genuine* indel evolutionary model. It should be noted here that the algorithm proposed here *assumes that the input MSA is correct*. Under this assumption, the algorithm approximately calculates the probabilities concerning the MSA.

### M1.1. Outline

Panel A of Figure 1 shows a flowchart of the procedures comprising our entire algorithm. Broadly speaking, the algorithm consists of three parts: (i) the “pre-processing” procedures that finally partition the entire input MSA into gapped segments and gapless segments separating them (steps ia-ic); (ii) enumerating the parsimonious local indel histories that can explain each gapped segment (step ii); and (iii) calculating the first-approximation of the augmented multiplication factor (Eq.(4.2.9c) of part I (Ezawa, Graur and Landan 2015a)) contributed from each gapped segment (step iii). The final results thus produced are put together, along with the overall factor (Eq.(4.2.9b) of part I), which is a function of the total length of the gapless segments, to provide the total occurrence probability of the entire MSA (Eq.(4.2.9a) of part I) as well as the relative probabilities among the parsimonious local indel histories that could explain each gapped segment (step iv). Panel B of Figure 1 schematically illustrates the procedures constituting the pre-processing part (steps ia-ic). The steps (ii) and (iii) will be described in Subsections M1.2 and M1.3, respectively.

After an input MSA is given [ Figure 1, step (o)], the algorithm first reduces the MSA to a binary pattern. In the binary pattern, each cell specified by a row (sequence) and a column (site) is given any of the two states: “presence” (denoted as “1”) when the cell is occupied by a residue, or “absence” (denoted as “0”) when it is filled with a gap [ step (ia)]. Then the algorithm decomposes the MSA into “gap-pattern block”s, or “block”s for short, each of which consists of contiguous columns with the same presence/absence pattern [ step (ib)]. Among such blocks, those containing no absence state play a distinct role as separators. If the MSA is correct, the existence of a gapless column indicates that no indel events occurred on or pierced through the column. This is a corollary of the phylogenetic correctness condition (e.g., Chindelevitch et al. 2006; Diallo et al. 2007), as explained in Subsection 3.4 of part I. Thus, gapless columns flanking a gapped segment *genuinely* delimit the indel events potentially responsible for the segment, *even in non-parsimonious histories*. Another note is that, although in general two contiguous gapless columns do not preclude indels in between them (also explained in Subsection 3.4 of part I), the algorithm described here ignores such indels between contiguous gapped columns, because it is only interested in parsimonious indel histories.

Then the algorithm makes a “cluster” out of a run of contiguous blocks containing the absence state and not separated from each other by gapless columns [ step (ic)]. Thus, each cluster spans between a gapless segment and the next gapless segment (or a MSA end). In this paper, we simply call such a cluster of gap-pattern blocks a “gapped segment.” As explained in Subsections 3.4 and 5.1 of part I, indel events and the probability of a local indel history in each gapped segment can be considered independently of events in the other gapped segments (even if we allow for non-parsimonious indel histories), as long as the indel model fulfills a set of conditions (as explained in Section 4 of part I).

Then, after the pre-processing part (step (i)) explained above, the two core parts follow: enumerating parsimonious local indel histories for each gapped segment (step (ii)), and calculating the occurrence probability of the segment (step (iii)). They will be explained in Subsections M1.2 and M1.3 below. The “post-processing” step (iv) will also be explained in Subsection M1.3.

### M1.2. Enumerating all parsimonious local indel histories

The first core part of our algorithm is itself an algorithm that attempts to enumerate all parsimonious local indel histories for each gapped segment. This core part consists of two subparts. (1) First it constructs an initial candidate for the local parsimonious indel histories, by identifying the unique Dollo parsimonious history (Farris 1977) for each gap-pattern block, and by merging together indel events of the same type in effectively contiguous blocks and along the same branch of the phylogenetic tree (Figure 2). (2) Then it iteratively searches for local indel histories whose events are fewer than or as many as those in the current candidate parsimonious histories (Figure 3), and it updates the set of candidate histories if such a history is found. It should be noted that, because our framework considers the input MSA to have resulted from an evolutionary process, the candidate indel histories must conform to the phylogenetic correctness condition (Chindelevitch et al. 2006; Diallo et al. 2007; see also Subsection 3.4 of part I (Ezawa, Graur and Landan 2015a)). We used the Dollo parsimonious state (Farris 1977) (in each gap-pattern block) as a starting point because it conforms to this condition. [The Dollo parsimony criterion (Farris 1977) seeks for an indel history consisting of the fewest events that can explain the gap-pattern, while only allowing for at most one insertion (per column or block) in order to keep the phylogenetic correctness.] In the following, we will explain these subparts in more detail.

(1) *Constructing an initial candidate of parsimonious local indel histories*. The first candidate history is constructed based on the block-wise Dollo parsimonious indel histories. The Dollo parsimonious indel history for each block can be easily and quickly constructed by a round-trip traversal of the (rooted) phylogenetic tree, first bottom-up and second top-down. In the bottom-up traversal, each node (*n*) is assigned the number of child nodes each of which has at least one extant descendant node with the “presence” state. Lets call the number *N*_*CDP*_(*n* ). When reaching the top (*i.e*., the root node *n*^*Root*^ ), the root is assigned the “presence” state if *N*_*CDP*_ (*n*^*Root*^) ≥ 2, otherwise it is assigned the “absence” state. Then, in the top-down traversal, each node (again *n*) is assigned the “presence” state, either if (a) *N*_*CDP*_ (*n*) ≥ 2, or if (b) *N*_*CDP*_ (*n*) = 1 and its parent is assigned the “presence” state. Otherwise, the node is assigned the “absence” state. Then, indels are inferred to have occurred only along the branches whose ends are assigned different “presence”/”absence” states.

Once the Dollo parsimony history is constructed for each block belonging to the gapped segment, the algorithm tries to reduce the number of indels by merging the effectively contiguous indel events of the same type (either all insertions or all deletions) and along the same branch in the sequence phylogeny (Figure 2). The “effectively contiguous” indel events can be either events in literally contiguous blocks (Figure 2, panel A) or events separated only by a (run of) block(s) that is (are) devoid of the “presence” state in any ‘downstream’ nodes (in the virtual temporal direction such that the event is viewed as a ‘deletion’) (panel B). When two events of the same type along the same branch are intervened by a block with the “presence” state in some ‘downstream’ nodes (the red “1” in panel C), however, the events are left unmerged. In most cases, this sub-part determines the unique parsimonious local indel history for each gapped segment.

(2) *Iteratively updating the set of candidate parsimonious local indel histories*. Not always and yet considerably frequently, the first sub-part doesn’t suffice to enumerate the parsimonious local indel histories. For example, in the situation illustrated in Figure 3, panel A, there could be another parsimonious history (panel C) on top of the initial history constructed in the first sub-part (panel B). In another example (panel D), there even exists a history (panel F) that requires less indel events than the intermediate candidate history (panel E), which requires as many indels as the initial, Dollo parsimonious history (not shown). Such histories can be found by iteratively updating the set of candidate parsimonious local indel histories, via “branch-and-merge” operations (panels G, H, F). A branch-and-merge operation is defined by first “branching” a ‘deletion’ event, that is, re-interpreting a ‘deletion’ event along a branch (panel G) as multiple independent ‘deletions’, each along one of the ‘child’ branches (panel H). Then, the “merging” process merges each resulting ‘deletion’ event with the effectively contiguous ‘deletion’ event(s), if at all, creating a new local indel history (panel F in this example). If the newly created history requires fewer indel events than the current candidate histories, then the new history replaces the current candidates. If the new scenario requires as many events as the current candidate(s), it joins the set of current candidates. Otherwise, the new history is discarded and, if some special conditions are met, the algorithm tries a more complex “branch-and-merge” operation as an attempt to exhaust all promising histories (detailed in Sub-subsection M1.2.2). For further details on this second sub-part, see Sub-subsections M1.2.1 and M1.2.2.

If you will, this second sub-part could be called a “**local multi-path downhill search algorithm.”** From each point, *i.e*., a local indel history, it examines only its neighborhoods, which are constructed by a single “branch-and-merge” operation. In this sense, it is a “**local** search.” Then, it keeps *only* those histories that consists of fewer indels than, or as few indels as, the current candidate. Thus it is “**downhill**.” At the same time, it keeps *all* histories that are found to have the same, “current-smallest,” number of indels. Hence it has the qualifier, “**multi-path**.”

#### M1.2.1. Assigning virtual temporal directions and ordering indel events

To exhaust (almost) all promising histories, the “branch-and-merge” processes (explained in Subsection M1.2, item (2)) are iterated from the “most influential” ‘deletion’ events, each of which ‘deletes’ a relevant sub-sequence from the largest number of aligned sequences, to the “least influential” ‘deletion’ events (Figure 4). Let us first explain what these single-quoted terms mean.

First, if the tree of the aligned sequences is rooted (panel A of Figure 4, left), it is converted to an unrooted tree (panel B), and all the indel events (panel A, right) are re-interpreted as ‘deletions’ (panel C). This could be done because the time direction could be arbitrarily assigned on an unrooted tree, and an insertion can be regarded as a deletion in the opposite time direction. The time direction may not be assigned consistently to all branches, for example when insertions and deletions coexist along a branch. However, this doesn’t matter and we will assign a unique ‘*virtual* time direction’ *to each indel event*, because it is only ‘deletion’ events with consistent directions that can be merged together, and because this re-interpretation is just a means to determine the order of the events that will go through the “branch-and-merge” operations.

Now, the ‘deletions’ will be sorted in descending order of the number of ‘deleted’ sequences (panel D), and will be processed from top to bottom of the list. The order is determined uniquely, except the ambiguity in the ordering among events that ‘delete’ the same number of sequences. This ambiguity is not expected to matter seriously, because the events that ‘delete’ the same number of sequences won’t be merged together in any “branch-and-merge” process.

A list of ‘deletions’ to be examined accompanies each candidate local indel history, and, each time a “branch-and-merge” operation is tried on the ‘deletion’ at the top of the list, the list is updated by removing the top ‘deletion’ just examined. If a “branch-and-merge” operation succeeds in finding a new promising candidate history, the new history is accompanied by a new list created by replacing the examined top ‘deletion’ with the resulting new ‘deletion(s).’ (The latter will be incorporated in the right order specified by the number of aligned sequences that the sub-sequence was ‘deleted from.’)

#### M1.2.2. Complex “branch-and-merge” operation for a special case

As explained in Subsection M1.2, item (2), our “local multi-path downhill search algorithm” first attempts a simple “branch-and-merge” operation on each ‘deletion’ event listed as described in Subsection M1.2.1. Now, assume that a simple branch-and-merge operation on a ‘deletion’ failed to produce a local history that requires no more indel events than the current candidate histories. Then, if two conditions are fulfilled, the algorithm will perform a complex “branch-and-merge” operation, which is described in this subsection. As explained in Subsection M1.2.1, the single quotation marks will indicate that the entity (or concept), especially that requiring a time-direction, is defined under the (virtual) time-direction in which the original indel event is interpreted as a ‘deletion.’

Here we clarify the two conditions required for a complex operation. (1) First of all, the original ‘deletion’ (like block *c* in panel A of Figure 5), say, along branch *e*_*o*_, must be effectively flanked on both sides within the same gapped segment, each side by a block with “presence” on some of the nodes ‘downstream’ of the branch *e*_*o*_, including its ‘lower-end’ node. (For example, in panel A of Figure 5, blocks *b* and *e* effectively flanks block *c*, but block *d* will be skipped.) (2) Then, ‘under’ the original ‘deletion,’ all branches that undergo ‘deletion’ events involving either of the “flanking” blocks are searched for (Figure 5 B and C, left) in a ‘bottom-up’ manner (Figure 6). Consider a situation where a set of ‘deletion’ events along such branches alone (Figure 5 B, left), or a set of such events plus an additional ‘deletion’ event (Figure 5 C, left), can ‘delete’ the subject block from all the sequences ‘affected’ by the original ‘deletion’ event in question. In this situation, the composite “branch-and-merge” operation can substitute the original ‘deletion’ event with a minimal set of such new ‘deletion’ events required to give the same effect on the aligned sequences (Figure 5 B and C, right). And each new ‘deletion’ event is merged with an old event involving at least one of effectively flanking blocks, if there is one along the same branch (Figure 5 B and C, right). And, in the block in question (*i.e.*, “block *c*” in Figure 5), from the ‘lower-end’ node of the original ‘deletion’ event to the ‘upper-end’ node of such new ‘deletion’ events, the “absence” state is flipped into the “presence” state (Figure 5 B and C, right). If no additional event is necessary, the new local indel history completely substitutes for the original one. If one additional event is necessary, the new local history joins the set of candidate histories. And, if more than one additional event are necessary (as in Figure 6 G), that is, unless condition (2) is fulfilled, the algorithm gives up “branch-and-merg”ing the original ‘deletion’ event.

A minimal set of ‘deletion’ events that substitute for the original event is searched for via a ‘bottom-up’ algorithm (Figure 6). Here the “minimal” set consists of the smallest number of events involving either or both of the effectively flanking blocks while accommodating the smallest number of additional events not involving either effectively flanking block. Actually, it is this algorithm that also examines how many additional events will be necessary to ‘delete’ the subject block(s) from all the sequences ‘affected’ by the original event. The algorithm starts with the external branches connecting to the sequences ‘affected’ by the original ‘deletion’ event, and ‘goes up’ the tree until reaching the branch (*e*_*o*_) along which the original ‘deletion’ event occurred (Figure 6, panel A). On each branch, the algorithm first examines whether there are ‘deletion’ events each of which involves at least one of the effectively flanking blocks (panel A). If there is at least one such effectively flanking ‘deletion’, the branch is labeled “directly absorbable”, and the algorithm assigns to it the branch itself (*e.g*., branches 1, 2, and 4 in panel B). Otherwise, the algorithm next examines whether there is a set of ‘downstream’ ‘deletion’ events, each involving at least one of the effectively flanking blocks, that jointly ‘delete’ the subject block from all the sequences ‘under’ the branch in question (*e.g.*, branch 5 in panel A). If there is such a set, the branch is labeled “indirectly absorbable”, and the algorithm assigns to it a set of “directly absorbable” branches ‘under’ it (*e.g.*, branch 5 in panel B). The set of “directly absorbable” branches ‘under’ an “indirectly absorbable” branch is the union of all the sets of “directly absorbable” branches assigned to the ‘child’ branches (panel C). If a branch is not labeled either as “directly absorbable” or “indirectly absorbable,” it is labeled “non-absorbable”, and no ‘descendant’ branch is assigned to it (panel D; *e.g*., branches 3 and 6 in panel B). Finally on branch *e*_*o*_, the algorithm counts the minimum number of additional ‘deletion’ events, that is, the number of its ‘child’ branches that are non-absorbable (panels E, F, G). The algorithm also constructs the minimum set of ‘deletion’ events that jointly substitute for the original ‘deletion’ (panels E, F, G). The minimum set is the union, over all the ‘child’ branches of branch *e*_*o*_, of the set of “directly absorbable” branches assigned to each ‘child’ branch (if it is “directly” or “indirectly” “absorbable”), or of the ‘child’ branch itself if it is “non-absorbable” (panel E, F, G). After this ‘bottom-up’ algorithm finishes, the minimum total number of additional events is given on branch *e*_*o*_, along which the original ‘deletion’ occurs.

To make the idea clearer, here we give a schematic code of the above ‘bottom-up’ algorithm. Let *e*_*o*_ denote the branch that underwent the original ‘deletion,’ *U*(*e*_*o*_) denote the set of all branches ‘under’ *e*_*o*_ (according to the (virtural) time direction consistent with that for the original ‘deletion’), *Ch*(*e*) denote the set of all ‘child’ branches of branch *e* (also according to the consistent (virtual) time direction), and *Da*(*e*) denote the set of “directly absorbable” branches assigned to branch *e*. The first recursive part is:

Foreach *e* ∈*U*(*e*_*o*_) (from the ‘lowest-level’ to the ‘highest-level’ ones (*i.e.*, those belonging to *Ch*(*e*_*o*_)))

If (*e* suffers a ‘deletion’ involving at least either of the effectively flanking blocks), then

*e* is “directly absorbable”, and

*Da*(*e*) = {*e*}.

Elseif (all *e*′ ∈ *Ch*(*e*) are either “directly” or “indirectly” “absorbable”), then

*e* is indirectly absorbable”, and

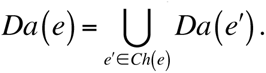

Else, then

*e* is “non-absorbable”, and

*Da*(*e*) is undefined.

End of if-elseif-else

End of foreach-loop.

Then, we give the subsequent part, which finishes the ‘bottom-up’ algorithm. Let *min*_*AD*_ (*e*_*o*_) denote the minimum number of additional ‘deletion’ events necessary to substitute for the original ‘deletion’ event along branch *e*_*o*_. And let *MS*_*SD*_ (*e*_*o*_) denote the minimum set of ‘deletion’ events that substitutes for the original event. Then we have the following schematic code:

Initially, *min*_*AD*_ (*e*_*o*_) = 0, and *MS*_*SD*_ (*e*_*o*_) is empty.

Foreach *e* ∈ *Ch*(*e*_*o*_ )

If (*e* is either “directly absorbable” or “indirectly absorbable”), then

*MS*_*SD*_ (*e*_*o*_) ← *MS*_*SD*_ (*e*_*o*_) ∪ *Da* (*e*).

Else (i.e. *e* is “non-absorbable”), then

*MS*_*SD*_ (*e*_*o*_) ← *MS*_*SD*_ (*e*_*o*_) ∪ {*e*}, and

*min*_*AD*_ (*e*_*o*_ )← *min*_*AD*_ (*e*_*o*_) + 1.

End of if-else

End of foreach-loop.

Here a left-pointing arrow represents that the entity on the right replaces that on the left. If *min*_*AD*_ (*e*_*o*_) = 0 (Figure 6, panel E), the newly constructed local indel history replaces the set of current candidate histories; if *min*_*AD*_ (*e*_*o*_) = 1 (panel F), the new history joins the set; if *min*_*AD*_ (*e*_*o*_) > 1 (panel G), the new history is discarded.

### M1.3. First-approximate calculation of absolute occurrence probability and relative probabilities

The second core part of our algorithm calculates the first approximation of the occurrence probability of a given entire MSA under a given phylogenetic tree and a given indel model, using only the contributions from parsimonious indel histories that are consistent with the MSA. The calculation is based on Eqs.(4.2.9a,b,c) of part I (Ezawa, Graur and Landan 2015a) for the probability of a given MSA, *α*[*s*_1_, *s*_2_,…, *s*_*N*^*X*^_]. The current version of the implementation of this core part only calculates the probability under Dawg’s indel model (Cartwright 2005), whose indel rate parameters (Eqs.(2.4.4a,b) of part I) are spatially and temporally homogeneous, and with a uniform length distribution of the ancestral sequence (*s*^*Root*^) at the root (*n*^*Root*^ ):

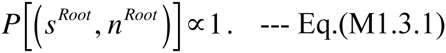

Thus, we always have 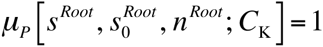 for every possible *s*^*Root*^ and for every potentially indel-accommodating region (*C*_K_ with ^∀^K ∈ {1,…, K_max_ } ). Here, as the “reference root state” 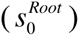, we do *not* use the concatenated root states of the block-wise Dollo parsimonious indel histories. Instead, as 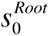, we use an array consisting solely of all sites corresponding to the gapless columns. Under a space-homogeneous model, this poses no problem. Let *N*_*GLC*_ (*α* [*s*_1_, *s*_2_,…, *s*_*N*^*X*^_]), or *N*_*GLC*_ for short, be the number of gapless columns in *α*[*s*_1_, *s*_2_,…, *s*_*N*^*X*^_]. Then, from Eq.(2.4.4c) of part I, we have:

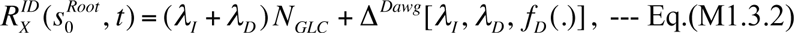

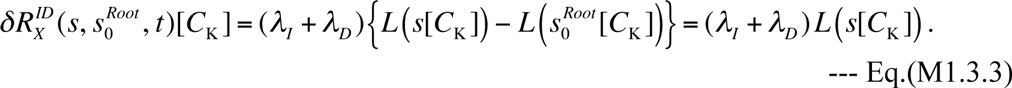

Here, *s*[*C*_K_] is the sub-sequence of the sequence *s* confined in the region *C*_K_, and we also used the fact that 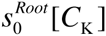 is always empty. Taking account of these equations, the “augmented multiplication factor” for the region *C*_K_ of the MSA *α*[*s*_1_, *s*_2_,…, *s*_*N*^*X*^_] given the tree *T*, whose specific expression is in Eq.(4.2.9c) of part I, is reduced to:

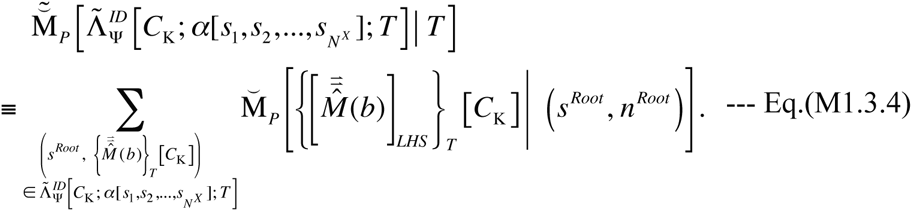

Here 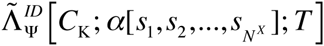 is the set of all indel histories along tree *T* (including the sequence state *s*^*Root*^ at the root node *n*^*Root*^) that can give rise to the portion of the MSA (*α*[*s*_1_, *s*_2_,…, *s*_*N*^*X*^_]) confined in the region *C*_K_. The summand of Eq.(M1.3.4) is given by a reduced form of Eq.(4.2.6b) of part I, that is,

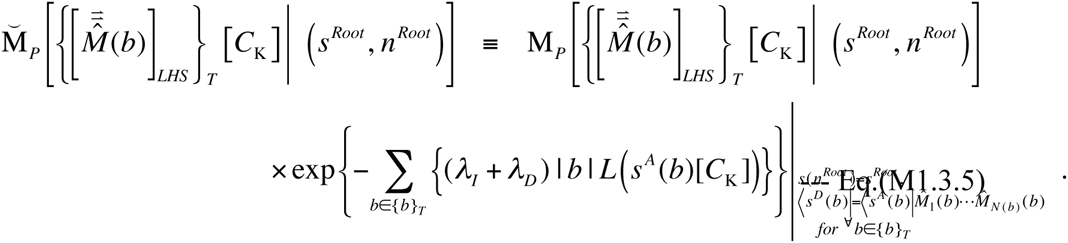

Here *s*^*A*^ (*b*) and *s*^*D*^ (*b*), respectively, are the sequence states at the ancestral and descendant nodes of branch *b*. And 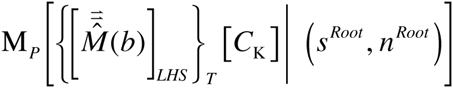 on the right hand side is the multiplication factor contributed from the local indel history that is the component of the local-history-set (LHS) equivalence class of indel histories along tree 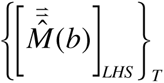, confined in the region *C*_K_, given the sequence state *s*^*Root*^ at the root. (Its specific expression is given by Eq.(4.2.4b) of part I.)

However, we cannot calculate the exact value of Eq.(M1.3.4), which collects the contributions from *all* local indel histories that could give rise to the portion of *α*[*s*_1_, *s*_2_,…, *s*_*N*^*X*^_] confined in *C*_K_. Instead, this second core part of our algorithm calculates its *first approximation* that is the summation of contributions from all *parsimonious* (*i.e.*, the fewest-indel) local indel histories potentially responsible for the MSA portion. Such an approximation can be given by a reduced form of Eq.(1.1.2b) of part II (Ezawa, Graur and Landan 2015b) with *N*_K_ = *N*_min_ [C_K_; *α*[*s*_1_, *s*_2_,…, *s*_*N*^*X*^_]; *T*] (or *N*min [*C*_K_] for short), *i.e*., the minimum number of indels necessary to give rise to the MSA portion. The first approximation is expressed as:

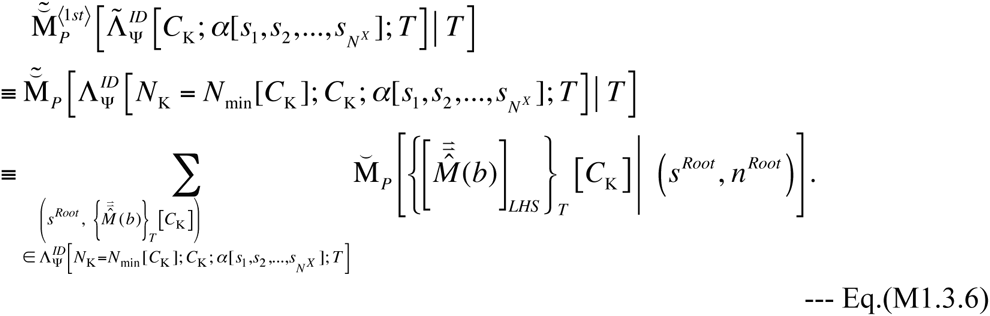

Using this, the first approximation of the probability of the entire MSA is given by a reduced form of Eqs.(4.2.9a,b) of part I. Their explicit expressions are:

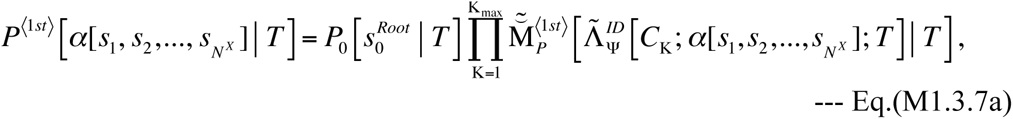

with

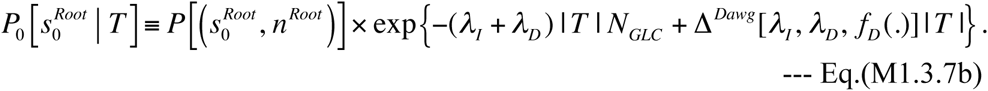

Here, 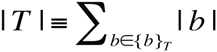 is the total length over all branches in the tree (*T* ). In general, the set of regions that can accommodate local indel histories, 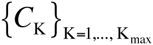, also contains the positions sandwiched by adjacent gapless columns within each single gapless segment. In the first approximation, the contribution, 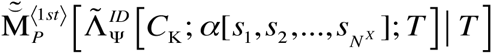, from each such sandwiched position is always trivial (*i.e*., unity). Thus, Eq.(M1.3.7a) could be further simplified as:

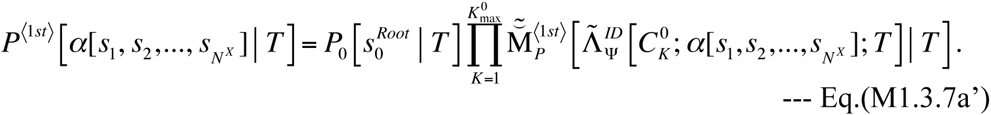

Here, 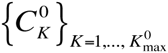 is the set of all gapped segments in the MSA. It is a subset of 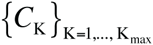, and thus 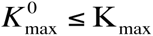 always holds. This Eq.(M1.3.7a’), supplemented by Eq.(M1.3.6) and Eq.(M1.3.7b), is the major output of the second core part, and of the entire algorithm.

As a byproduct, the second core part also outputs the relative probabilities among the parsimonious local indel histories that can give rise to the same local MSA confined in each gapped segment, 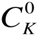. The relative probability of each parsimonious local history, 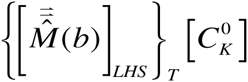, is calculated as:

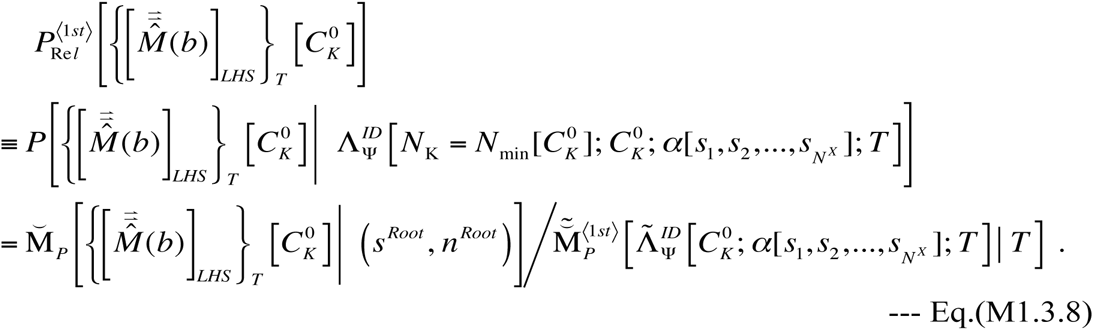

### M2. Validation analyses

In order to validate our algorithm described in Section M1, we conducted various analyses by applying the algorithm or its components to the data synthesized either manually or via simulations. (The algorithm, including its components, was implemented in Perl and is available at the FTP repository of the Bioinformatics Organization (Ezawa 2013).)

### M2.1. Manual validation of parsimony component of our algorithm

To validate the first core component, *i.e*., an algorithm that attempts to enumerate all parsimonious local indel histories for each gapped segment (described in Subsection M1.2), we first applied it to an extensive set of manually synthesized data. The manual data set consists of mini-MSAs with various gap-configurations and with various phylogenetic relationships between the sequences, as shown in panels A-R of Figure 7. Then, we manually inspected the outputs (illustrated in Figures 8-25), and confirmed that the algorithm indeed exhausts all possible parsimonious local indel histories for most of the inputs (panels A-Q of Figure 7).

A caveat is that this first implementation of the algorithm may miss some parsimonious local histories occasionally, for example for the gap-configuration in panel R of Figure 7. Our current implementation did find a parsimonious history that can result in this configuration (Figure 25). However, this is not all; the gap-configuration in Figure 7 R is of an “intersection between cousins” type according to the classification by Fredslund et al. (2003), and can be explained also by another parsimonious history (Figure 26). Because our “branch-and-merge” operation cannot reach this type of local indel histories, we need to introduce another operation to find them. Good news is that such gap-configurations are expected to be very rare because these histories have probabilities at least of the 4th order in the indel rate. Another caveat is on the gap-configuration in Figure 7 K. Although our algorithm output Figure 18 as a “parsimonious” local indel history, it is actually unlikely to give rise to the pattern, at least via Dawg (Cartwright 2005). If a *genuine* molecular evolution simulator like Dawg creates the pattern, it will probably be through histories shown in Figure 27, although they require more indel events than the history in Figure 18. This is related with the issue of a three-event indel history discussed in Subsection 3.3 of part I (Ezawa, Graur and Landan 2015a). These issues will also be addressed in the future version of the implementation.

### M2.2. Simulations to prepare input MSA sets

To validate the entire algorithm described in Section M1, we prepared three sets of MSAs using the *genuine* sequence evolution simulator, Dawg (Cartwright 2005). We performed all simulations using the same Zipf power-law distribution, 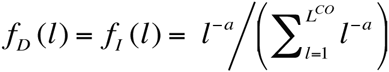, with the exponent *a* = 1.6 and the cut-off indel length 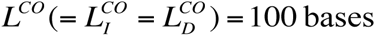. The exponent *a* = 1.6 is typical among empirically observed values (Gonnet et al. 1992; Gu and Li 1995; Zhang and Gerstein 2003; Chang and Benner 2004; Fan et al. 2007; Cartwright 2009). The cut-off length was chosen in order to prevent it from taking extremely long to search for parsimonious local indel histories; with our current implementation, the search could be very long when a gapped segment contains at least one long gap (see Discussion D1.1 for a possible solution). Each of the simulations started with a random ancestral DNA sequence that is 1,000 bases long. In each simulation, we labeled all the internal nodes of the input tree, in order to keep the ancestral sequences aligned with the “extant” sequences (at the external nodes). Other parameters and options were set at default values unless otherwise stated. We created three input MSA sets, 1A, 1B, and 2.

**Set 1A** consists of 100,000 MSAs, each of which was simulated along a 3-taxon tree starting at a root with three children, and the lengths of the three branches were all set at 0.05 (substitutions per base). The total rates of insertions and deletions were set at *λ*_*I*_ = *λ*_*D*_ = 0.1 (per expected substitution), which are close to the upper-bounds for neutrally evolving mammalian DNA sequences (Lunter 2007; Cartwright 2009).

**Set 1B** is similar to Set 1A, expect that all branch lengths were set at 0.2 (substitutions per base).

We prepared these two sets, 1A and 1B, because validating the theoretically predicted occurrence probabilities of local gap configurations necessitated a large number of MSAs simulated under identical parameter settings.

**Set 2** consists of 9,900 MSAs, each of which was simulated along a 16-taxon tree. It is actually a union of 33 subsets, each consisting of 300 MSAs simulated under a same parameter setting (the topology and branch lengths of the phylogenetic tree, and a value of the indel rates, *λ*_*I*_ = *λ*_*D*_ ). These parameter settings were chosen to represent typical values for the MSAs in a benchmark database, BAliBASE (Thompson et al. 2005). More precisely, the parameters were chosen to reproduce the 5th, 10th, 25th, 50th, 75th, 90th, and 95th percentiles of each attribute. In each subset, one was varied out of the following 5 attributes, namely, the mean branch length, the gap content, the coefficient of variation (CV = standard deviation / mean) of the branch length, the CV of the number of branches separating a pair of leaves, and the CV of the distance from the root to a leaf. The remaining 4 out of the 5 attributes were kept at the median (*i,e.*, the 50th percentile) for each subset. After removing the redundancies out of the 35 parameter settings, 33 were left (for trees and indel rates, see Figure 28). This Set 2 is therefore expected to reproduce various properties of the real-life MSAs.

The control files used to generate these simulated datasets, including the phylogenetic trees and indel model parameters, are available in the LOLIPOG package at the FTP repository of the Bioinformatics Organization (Ezawa 2013).

### M2.3. Comparing parsimonious local indel histories with true history

Rigorously speaking, unless the record is kept on the true local indel history that created each observed gapped segment, we cannot compare it with the predicted (parsimonious) histories. Nevertheless, if we have the ancestral sequence states at all the internal nodes aligned with the “extant” sequences at the external nodes, we can *approximately* judge whether the true local indel history matches one of the predicted (parsimonious) histories, by comparing the gap states of all the true ancestral sequences (in the segment in question) with the ancestral gap states in each predicted history. [It should be noted, however, that the judgment is only approximately correct, because the same set of ancestral sequence states could result from more than one local indel history if non-parsimonious histories are also counted in.] Because Dawg can output the alignment of ancestral sequences at the labeled internal nodes with the “extant” sequences at the external nodes, we took advantage of this function and examined whether the true ancestral gap states in each instance of a gapped segment match those predicted by one of the parsimonious local indel histories. If there is a match, we registered the instance as “parsimonious,” and recorded which parsimonious history can produce the true ancestral gap states; otherwise, we registered the instance as “non-parsimonious.” Dawg sometimes creates gapped segments containing null columns, in each of which all extant sequences are occupied by gaps. Because indel histories that create null columns can never be parsimonious in the sense of MSA reconstruction, we also registered gapped segments containing such null columns as “non-parsimonious.”

### M2.4. Correlation analysis to validate predicted absolute occurrence probabilities of gapped segments

To examine whether or not our first approximation of the augmented multiplication factor, Eq.(M1.3.6), works well, we first counted instances of gapped segments that occurred in each of the simulated sets A1 and A2 without reaching either MSA end, and that showed a particular gap-configuration, say, *G*_*a*_. Then we compared the count of instances (*i.e*., the absolute frequency) of gap-configuration *G*_*a*_ with its theoretical prediction, 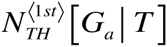, which was calculated using Eq.(M1.3.6) (for a gapped segment *C*_K_ that exhibits the gap-configuration *G*_*a*_) as:

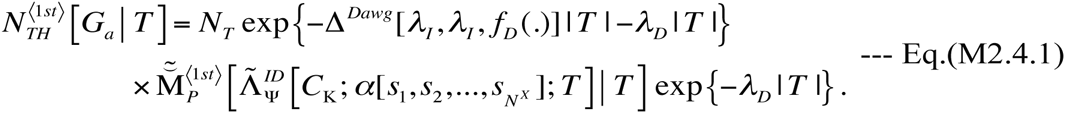

Here *N*_*T*_ is the total number of sites in the root sequences where insertions/deletions potentially occur. And, roughly speaking, exp {−Δ^*Dawg*^[*λ*_*I*_, *λ*_*I*_, *f*_*D*_ (.)] | *T* | − *λ*_*D*_ | *T* |} is the probability that the left-flanking gapless column remained undeleted, and exp {−*λ*_*D*_ | *T* |} is the conditional probability that the right-flanking gapless column remained undeleted given the gapped segment and the left-flanking gapless column. In this study, we simply set *N*_*T*_ = 100, 000 × 1, 000 = 10^8^, which is the total number of bases in the root ancestral sequences in each input set. We used only those gap-configurations each of which is expected to occur 5 or more times in each dataset.

We compared the absolute frequencies of the gap-configurations predicted by Eq.(M2.4.1) against their actual frequencies in each simulated dataset by performing the correlation and linear regression analyses between their square roots. We did so based on the following rationale. The count of each gap-configuration in each simulated dataset is expected to roughly follow a Poison distribution, in which the standard error of the count of events is the square root of its mean. Therefore, the square root of the simulated count is expected to have a standard error that is roughly uniform independently of the gap-configuration. This uniformity of the standard error is a major assumption underlying the correlation and linear regression analyses.

Before the analyses in this and the next subsections, we pre-processed the simulated MSAs in order to avoid complexities caused by the equivalence relations (Eqs.(A1.3c,d,c’,d’) in Appendix A1 of part I (Ezawa, Graur and Landan 2015a)), as explained in Subsection 3.3 of part I, and by similar equivalence relations regarding independent insertions. Specifically, we first removed null columns. Then we swapped two adjacent gap-pattern blocks when they satisfied the following two conditions. (1) A block shows the “presence” state only in sequences that show the “absence” state in the other block. (2) In the MSA, the highest sequence with the “presence” state in the left block is higher than that in the right block.

### M2.5. Correlation analysis to validate predicted relative probabilities among parsimonious local indel histories

To examine whether or not our formula for the relative probability, Eq.(M1.3.8), works well with each of the simulated sets, A1, A2, and B, we first calculated Eq.(M1.3.8) for all alternative parsimonious local indel histories of all “parsimonious” instances of gapped segments that do not reach either MSA end. Then, we distributed the histories enumerated for each input set into 20 non-overlapping bins of 5% width that jointly span the open interval, (0, 1), of the theoretical relative probability. (The histories with the relative probability = 1 were excluded from the analyses because they could cause the performance to be unfairly overrated.) In each bin, we counted the total number of instances of alternative histories considered as well as the number of actual instances of “correct” histories, whose ancestral gap states matched the true ones. Then, we compared the simulated proportion (*i.e.*, relative frequency) of “correct” histories in each bin with the theoretically predicted probability that the history is “correct,” 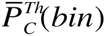. The 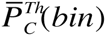 for each bin was calculated by averaging the relative probabilities given by Eq.(M1.3.8) over all instances of the considered alternative histories in the bin. We performed the correlation and linear regression analyses, using the actual relative frequency in each simulated dataset as the explanatory variable, and using the predicted probability as the response variable. For the analyses, we used averages weighted by the reciprocal of the variance of the predicted probability, *i.e*., 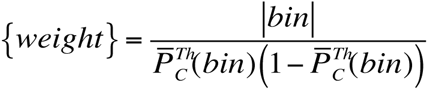, where |*bin*| is the total number of instances of considered alternative histories in the bin.

Similar correlation and linear regression analyses were conducted also on the most likely (ML) parsimonious local indel histories alone, as well as on the least likely (LL) parsimonious histories alone.

## Authors’ contributions

KE conceived of and mathematically formulated the theoretical framework in this paper, implemented the key algorithms, participated in designing the study, performed all the analyses of the simulated MSA datasets, and drafted the manuscript. DG and GL participated in designing the study, helped with the interpretation of the data, and helped with the drafting of the manuscript. GL also determined the parameter sets for set 2 of simulated MSAs by analyzing the benchmark MSA dataset from BAliBASE.

## Acknowledgements

This study is dedicated to the late Dr. Keiji Kikkawa, who was a renowned theoretical physicist, one of the key pioneers of the string field theory of the elementary particle physics, and the best ever mentor of K.E. We are grateful to Dr. R. A. Cartwright at Arizona State University for his useful information and discussions that inspired this study. We appreciate the logistic support and the feedback of Dr. Tetsushi Yada at the Kyushu Institute of Technology. We would also like to thank the three anonymous referees of the predecessor manuscript entitled: “Framework that enables approximate lilelihood analysis of insertions/deletions on multiple sequence alignment.” Their comments helped drastically improve the study itself, not to mention the manuscript. This work was a part of the project, “Error Correction in Multiple Sequence Alignments,” which was funded by US National Library of Medicine [grant number LM010009-01 to Dan Graur and Giddy Landan at the University of Houston]. The later stage of this work was also supported by Grants-in-Aid No. 221S0002, which was awarded to Tetsushi Yada by the Ministry of Education, Culture, Sports, Science and Technology of Japan.

## References

Bishop MJ, Thompson EA. 1986. Maximum likelihood alignment of DNA sequences. J. Mol Biol. 190:159–165.

Bouchard-Côté A, Jordan MI. 2013. Evolutionary inference via the Poisson indel process.s Proc Natl Acad Sci USA. 110:1160–1166.

Bradley RK, Holmes I. 2007. Transducers: an emerging probabilistic framework for modeling indels on trees. Bioinformatics 23:3258–3262.

Britten RJ. 2002. Divergence between samples of chimpanzee and human DNA sequences is 5%, counting indels. Proc. Natl. Acad. Sci. USA 99:13633–13635.

Britten RJ, Rowen L, Willians J, Cameron RA. 2003. Majority of divergence between closely related DNA samples is due to indels. Proc. Natl. Acad. Sci. USA 100:46614665.

Cartwright RA. 2005. DNA assembly with gap (Dawg): simulating sequence evolution. Bioinformatics 21:iii31–iii38.

Cartwright RA. 2009. Problems and solutions for estimating indel rates and length distribution. Mol Biol Evol. 26:473–480.

Chang MSS, Benner SA. 2004. Empirical analysis of protein insertions and deletions determining parameters for the correct placement of gaps in protein sequence alignments. J Mol Biol. 341:617–631.

Chindelevitch L. Li Z, Blais E, Blanchette M. 2006. On the inference of parsimonious evolutionary scenarios. J Bioinform Comput Biol. 4:721–744.

Diallo AB, Makarenkov V, Blanchette M. 2007. Exact and heuristic algorithms for the indel maximum likelihood problem. J Comput Biol. 14:446–461.

Ezawa K. 2013. LOLIPOG: Log-likelihood for the pattern of gaps in MSA. [http:///www.bioinformatics.org/ftp/pub/lolipog/]

Ezawa K, Graur D, Landan G. 2015a. Perturbative formulation of general continuous-time Markov model of sequence evolution via insertions/deletions, Part I: Theoretical basis. bioRxiv doi: http://dx.doi.org/10.1101/023598.

Ezawa K, Graur D, Landan G. 2015b. Perturbative formulation of general continuous-time Markov model of sequence evolution via insertions/deletions, Part II: Perturbation analyses. bioRxiv doi: http://dx.doi.org/10.1101/023606.

Ezawa K, Graur D, Landan G. 2015c. Perturbative formulation of general continuous-time Markov model of sequence evolution via insertions/deletions, Part IV: Incorporation of substitutions and other mutations. bioRxiv doi: http://dx.doi.org/10.1101/023622.

Fan Y, Wang W, Ma G, Liang L, Shi Q, Tao S. 2007. Patterns of insertion and deletion in mammalian genomes. Curr Genomics 8:370–378.

Farris JS. 1977. Phylogenetic analysis under Dollo’s law. Syst Zool. 26:77–88

Felsenstein J. 1981. Evolutionary trees from DNA sequences: a maximum likelihood approach. J Mol Evol. 17:368–376.

Felsenstein J. 2004. Inferring Phylogenies. Sunderland (MA), Sinauer Associates.

Fredslund J, Hein J, Scharling T. 2003. A large version of the small parsimony problem. In: Benson G, Page R, editors. WABI 2003, LNBI 2812. Heidelberg, Springer-Verlag. p. 417–432.

Gascuel O (editor). 2005. Mathematics of Evolution and Phylogeny. New York, Oxford University Press.

Gonnet GH, Cohen MA, Benner SA. 1992. Exhaustive matching of the entire protein sequence database. Science 256:1443–1445.

Graur D, Li WH. 2000. Fundamentals of Molecular Evolution, 2nd ed. Sunderland (MA), Sinauer Associates.

Gu X, Li WH. 1995. The size distribution of insertions and deletions in human and rodent pseudogenes suggests the logarithmic gap penalty for sequence alignment. J Mol Evol. 40:464–473.

Holmes I, Bruno WJ. 2001. Evolutionary HMMs: a Bayesian approach to multiple sequence alignment. Bioinformatics 17:803–820.

Kent WJ, Baertsch R, Hinrichs A, Miller W, and Haussler D. 2003. Evolution’s cauldron: duplication, deletion, and rearrangement in the mouse and human genomes. Proc Natl Acad Sci USA 100:11484–11489.

Landan G, Graur D. 2009. Characterization of pairwise and multiple sequence alignment errors. Gene 441:141–147.

Löytynoja A, Goldman N. 2008. Phylogeny-aware gap placement prevents errors in sequence alignment and evolutionary analysis. Science 320:1632–1635.

Lunter G. 2007. Probabilistic whole-genome alignments reveal high indel rates in the human and mouse genomes. Bioinformatics 23:i289–i296.

Lunter GA, Miklós I, Drummond A, Jensen JL, Hein J. 2005. Bayesian coestimation of phylogeny and sequence alignment. BMC Bioinformatics 6:83.

Lunter G, Rocco A, Mimouni N, Heger A, Caldeira A, Hein J. 2008. Uncertainty in homology inferences: assessing and improving genomic sequence alignment. Genome Res. 18:298–309.

Lynch M. 2007. The Origins of Genome Architecture. Sunderland (MA), Sinauer Associates.

Miklós I, Lunter GA, Holmes I. 2004. A “long indel” model for evolutionary sequence alignment. Mol Biol Evol. 21:529–540.

Miklós I, Novák á, Satija R, Lyngsø R, Hein J. 2009. Stochastic models of sequence evolution including insertion-deletion events. Stat Methods Med Res. 18:453–485.

Novák á, Miklós I, Lyngsø R, Hein J. 2008. StatAlign: an extendable software package for join Bayesian estimation of alignments and evolutionary trees. Bioinformatics 24:2403–2404.

Paten B, Herrero J, Fitzgerald S, Beal K, Flicek P, Holmes I, Birney E. 2008. Genomewide nucleotide-level mammalian ancestor reconstruction. Genome Res. 18:18291843.

Redelings BD, Suchard MA. 2005. Joint Bayesian estimation of alignment and phylogeny. Syst Biol. 54:401–418.

Redelings BD, Suchard MA. 2007. Incorporating indel information into phylogeny estimation for rapidly emerging pathogens. BMC Evol Biol. 7:40.

Rivas E. 2005. Evolutionary models for insertions and deletions in a probabilistic modeling framework. BMC Bioinformatics 6:63.

Rivas E, Eddy SR. 2008. Probabilistic phylogenetic inference with insertions and deletions. PLoS Comput Biol. 4:e1000172.

Strope CL, Abel K, Scott SD, Moriyama EN. 2009. Biological sequence simulation for testing complex evolutionary hypothesis: indel-Seq-Gen version 2.0. Mol Biol Evol. 26:2581–2593.

Suchard MA, Redelings BD. 2006. BAli-Phy: simultaneous Bayesian inference of alignment and phylogeny. Bioinformatics 22:2047–2048.

The Chimpanzee Sequencing and Analysis Consortium. 2005. Initial sequence of the chimpanzee genome and comparison with the human genome. Nature 437:69–87.

The International Chimpanzee Chromosome 22 Consotrium. 2004. DNA sequence and comparative analysis of chimpanzee chromosome 22. Nature 429:382–388.

Thompson JD, Koehl P, Ripp R, Poch O. 2005. BAliBASE 3.0: Latest development of the multiple sequence alignment benchark. PROTEINS 61:127–136.

Thorne JL, Kishino H, Felsenstein J. 1991. An evolutionary model for maximum likelihood alignment of DNA sequences. J Mol Evol. 33:114–124.

Westesson O, Lunter G, Paten B, Holmes I. 2012. Accurate reconstruction of insertion-deletion histories by statistical phylogenetics. PLoS One 7:e34572.

Yang Z. 2006. Computational Molecular Evolution. New York (NY), Oxford University Press.

Zhang ZL, Gerstein M. 2003. Patterns of nucleotide substitution, insertion and deletion in the human genome inferred from pseudogenes. Nucleic Acids Res. 31:5338–5348.

